# Identification of leukemic and pre-leukemic stem cells by clonal tracking from single-cell transcriptomics

**DOI:** 10.1101/500108

**Authors:** Lars Velten, Benjamin A. Story, Pablo Hernández-Malmierca, Simon Raffel, Daniel R. Leonce, Jennifer Milbank, Malte Paulsen, Aykut Demir, Chelsea Szu-Tu, Robert Frömel, Christoph Lutz, Daniel Nowak, Johann-Christoph Jann, Caroline Pabst, Tobias Boch, Wolf-Karsten Hofmann, Carsten Müller-Tidow, Andreas Trumpp, Simon Haas, Lars M. Steinmetz

**Affiliations:** Centre for Genomic Regulation (CRG), The Barcelona Institute of Science and Technology, Dr. Aiguader 88, Barcelona 08003, Spain; Universitat Pompeu Fabra (UPF), Barcelona, Spain; European Molecular Biology Laboratory (EMBL), Genome Biology Unit, 69117 Heidelberg, Germany; Swiss Federal Institute of Technology (ETH) Zurich, Department of Biosystems Science and Engineering, 4058 Basel, Switzerland; Heidelberg Institute for Stem Cell Technology and Experimental Medicine (HI-STEM gGmbH), 69120 Heidelberg, Germany; Division of Stem Cells and Cancer, Deutsches Krebsforschungszentrum (DKFZ) and DKFZ-ZMBH Alliance, 69120 Heidelberg, Germany; Department of Internal Medicine V, Hematology, Oncology and Rheumatology, University of Heidelberg, 69120 Heidelberg, Germany; European Molecular Biology Laboratory (EMBL), Flow Cytometry Core Facility, 69117 Heidelberg, Germany; Department of Hematology and Oncology, Medical Faculty Mannheim, Heidelberg University, 68167 Mannheim, Germany; Molecular Medicine Partnership Unit (MMPU), University of Heidelberg and European Molecular Biology Laboratory (EMBL), Heidelberg, Germany; German Cancer Consortium (DKTK), 69120 Heidelberg, Germany; Berlin Institute of Health (BIH), 10178 Berlin, Germany; Charité-Universitätsmedizin, 10117 Berlin, Germany; Berlin Institute for Medical Systems Biology, Max Delbrück Center for Molecular Medicine in the Helmholtz Association, 10115 Berlin, Germany; Department of Genetics, Stanford University School of Medicine, Stanford, California 94305, USA; Stanford Genome Technology Center, Palo Alto, California 94304, USA

**Keywords:** Acute Myeloid Leukemia, Cancer evolution, Cancer stem cells, Single-cell genomics

## Abstract

Cancer stem cells drive disease progression and relapse in many types of cancer. Despite this, a thorough characterization of these cells remains elusive and with it the ability to eradicate cancer at its source. In acute myeloid leukemia (AML), leukemic stem cells (LSCs) underlie mortality but are difficult to isolate due to their low abundance and high similarity to healthy hematopoietic stem cells (HSCs). Here, we demonstrate that LSCs, HSCs, and pre-leukemic stem cells can be identified and molecularly profiled by combining single-cell transcriptomics with lineage tracing using both nuclear and mitochondrial somatic variants. While mutational status discriminates between healthy and cancerous cells, gene expression distinguishes stem cells and progenitor cell populations. Our approach enables the identification of LSC-specific gene expression programs and the characterization of differentiation blocks induced by leukemic mutations. Taken together, we demonstrate the power of single-cell multi-omic approaches in characterizing cancer stem cells.

## Introduction

Tissues with high cellular turnover, such as the hematopoietic system or the intestine, depend on ‘professional’ adult stem cells for their continuous regeneration^1^. Oncogenic mutations in these cells can cause cancers that maintain a hierarchical organization reminiscent of the tissue of origin. Only cancer stem cells (CSCs), residing at the top of the hierarchy, are able to fuel long-term cancer growth and drive relapse, whereas the bulk of the cancer consists of rapidly dividing cells with limited capacity for self-renewal, i.e. cells that exhaust their replicative potential after a finite number of divisions^2–4^. Due to their stem cell-like properties, CSCs constitute an important driver of relapse, but their low division rates make them difficult to target therapeutically. Tools that permit the confident identification and characterization of CSCs are therefore urgently needed.

Acute myeloid leukemia (AML) serves as a paradigm for the study of cancer stem cells^5^. In 10-20% of healthy individuals over age 70, the acquisition of pre-leukemic mutations in hematopoietic stem cells (HSCs) results in the dominance of a small number of HSC-derived clones, a process termed Clonal Hematopoiesis of Indeterminate Potential (CHIP)^6,7^. While such pre-leukemic stem cells (pre-LSCs) are capable of giving rise to healthy blood and immune cells, additional mutations can cause a complete block in differentiation and thereby result in the malignant expansion of aberrant progenitor cells^8^. The accumulation of these so-called ‘blast’ cells is ultimately fueled by the presence of leukemic stem cells (LSCs). Classic chemotherapy regimens primarily target actively cycling ‘blast’ cells and initially lead to remission. Since quiescent or protected LSCs often avoid eradication, relapse rates are high with 5-year survival rates below 15% for patients over the age of 60. A key goal is therefore to identify therapeutic strategies for targeting LSCs, while sparing healthy HSCs^9–11^. Characterizing gene expression differences between HSCs, pre-LSCs and LSCs would be a valuable step towards that goal.

Previously, LSC-specific gene expression patterns were characterized by isolating cells positive for stem cell-specific surface markers used in the healthy hematopoietic system, such as CD34^12–15^. More recently, leukemic engraftment rates in xenotransplant models were correlated with gene expression^16–19^. However, these approaches all measure impure populations of cells. Here, we propose that by measuring mutational status and gene expression in single cells simultaneously, cancer stem cells can be uniquely distinguished from both mature cancer cells (based on gene expression) and healthy stem cells (based on mutational status) (Figure 1a). Finally, pre-LSCs are thought to typically carry mutations associated with CHIP (e.g. *Dnmt3a*) but not mutations associated with leukemia (e.g. *Npm1*)^7,20,21^, potentially enabling their identification by profiling both known leukemic and known preleukemic mutations.

**Figure 1.**
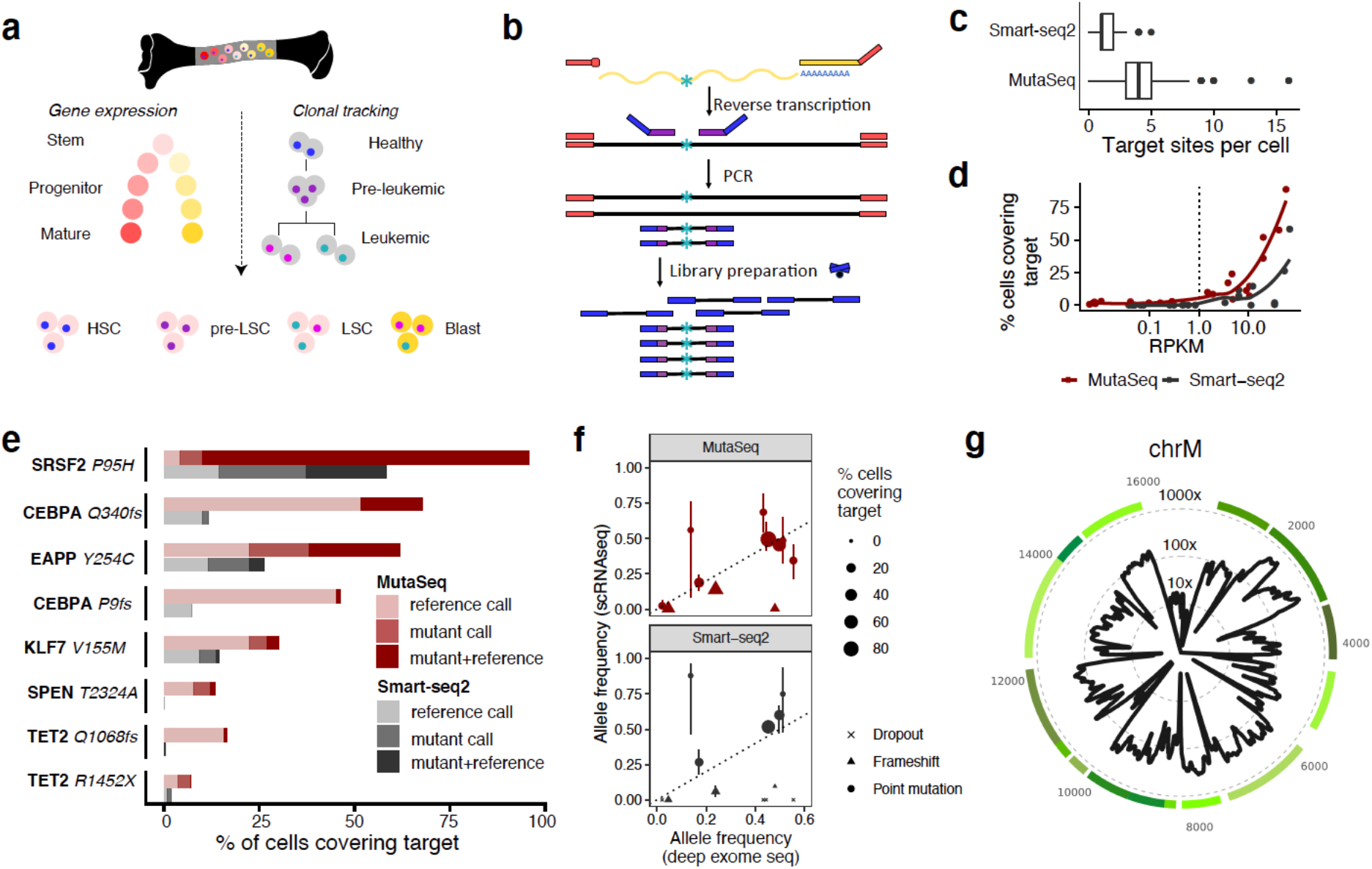
MutaSeq for high-quality single-cell RNA-seq data with clonal information. See also Figure S1. a. Overview of the study. HSC, Hematopoietic stem cell; (pre-)LSC, (pre#x002D;)leukemic stem cell; Blast, mature leukemic blast. b. Overview of the MutaSeq method. Targeting primers (purple) are included during the cDNA amplification step of the Smart-seq2 protocol. Targeting primers are directly fused to illumina library adapters (blue) and therefore get amplified efficiently during library preparation. Tagmentation introduces the same adapters to the full-length cDNA product. c. Number of target sites covered per cell, across n=206 (Smart-seq2) or n=658 CD34+ (MutaSeq) bone marrow cells from patient P1. See Methods section on ‘Data Visualization’ for a definition of boxplot elements. d. Mean gene expression of genes containing mutations of interest is plotted against the fraction of cells in which the mutation is covered. e. Fractions of cells covering key non-synonymous mutations observed in the patient. Reference call: The reference allele was observed. Mutant call: the mutant allele, as defined by bulk exome sequencing (Table S1) was observed. Mutant+Reference: both alleles were observed. f. Allele frequency estimates derived from deep exome sequencing compared to allele frequency estimates derived from MutaSeq (red dots) or Smart-seq2 (grey dots). Dot size indicates coverage at target site. Point shape indicates the type of mutation. g. Coverage of the mitochondrial genome in MutaSeq data. See Figure S1j for a comparison across methods. Green line segments correspond to genes in the mitochondrial genome.

While we and others have demonstrated the utility of single-cell genomics for mapping hematopoietic differentiation hierarchies^22–24^, tracking mutations or clones in single-cell gene expression data remains difficult. Previous work has amplified somatic mutations from cDNA^25–28^, extracted mutational information from single-cell RNA-seq reads^29^, or processed both genomic DNA and RNA from single cells^30–32^. However, these protocols suffer from a lack of confidence in assigning cells to clones and/or require prior knowledge of genomic mutation sites. As an alternative tool for clonal tracking, the use of endogenous mitochondrial mutations as clonal markers has been proposed, obviating the need for prior knowledge of genomic mutations^33,34^. However, the application of these methods to characterize LSCs has not been demonstrated, and in particular requires the ability to reliably detect clonal expansion events, associate clinically-relevant coding mutations to clones with high confidence, and draw statements on gene expression changes between clones.

Here, we introduce MutaSeq, a workflow that amplifies nuclear mutations from cDNA, and *mitoClone*, a computational tool that achieves high-confidence clonal assignments and de novo discovery of clones using mitochondrial marker mutations when available. MutaSeq data from four AML patients allowed us to distinguish HSCs, pre-LSCs, LSCs, and progenitor/blast populations. Thereby, we identified transcriptomic consequences of leukemic and pre-leukemic mutations relevant to stem cells. Additionally, we characterized the contribution of different leukemic and pre-leukemic clones to healthy and disease-specific bone marrow populations with unprecedented detail. Together, our results demonstrate cancer stem cell identification and characterization by simultaneous mapping of genomic and mitochondrial mutations in single-cell transcriptomes.

## Results

### MutaSeq provides high coverage of genomic and mitochondrial mutations

To establish a robust experimental setup for the clonal tracking of human cells in single-cell transcriptomic data, we evaluated various modifications of the Smart-seq2 protocol aimed at increasing coverage at polymorphic genomic sites of interest (Figure S1a). We found that inclusion of targeting primers during reverse transcription frequently resulted in the formation of undesired byproducts, especially when targeting a higher number of sites (Figure S1a-d). In contrast, when sites of interest were targeted during cDNA amplification, we obtained high quality transcriptome data while increasing the average number of target sites captured per cell by 2-4-fold compared to a non-targeted approach (Figures 1b,c, and S1a,b). An automated pipeline for primer design that minimizes off-target sites and potential primer-dimer formation is available at https://github.com/veltenlab/PrimerDesign (see also Methods section). MutaSeq stably works with up to 30-40 primer pairs (targets); when higher numbers of primers are included, library quality progressively decreases (Figure S1e). In a test with only highly expressed target genes, target amplicons are created from all primer pairs in virtually all single cells (Figure S1f,g).

To evaluate MutaSeq, we performed deep exome sequencing of an AML patient (P1) and designed primers targeting 14 nuclear mutations (Table S1, S5). We then systematically compared the performance of MutaSeq and non-targeted Smart-seq2 on CD34+ cells from this patient. MutaSeq increased the number of target sites covered per single cell from a median of 1 to a median of 4 (Figure 1c-e) and recapitulated the variant allele frequencies estimated by exome sequencing with higher accuracy than Smart-seq2 (Figure 1f), while maintaining comparable transcriptome data quality (Figure S1h,i). Both methods underestimated the abundance of frameshift mutations, possibly as a consequence of nonsense mediated decay^35^ (Figure 1e,f). While our data do not provide statistical evidence for an effect of target gene length or sequence complexity on dropout, we cannot exclude such effects.

Importantly, both methods provide an excellent coverage of the mitochondrial genome (mean ~100X mitochondrial coverage given a mean sequencing depth of ~788,000 total reads per cell, Figure 1g), unlike most other single-cell RNA-seq protocols applied in the context of clonal tracking^27–29–31^ (Figure S1j). Together, these results demonstrate that MutaSeq efficiently covers the mitochondrial genome in single-cell RNA-sequencing experiments and provides improved coverage of genomic target sites compared to Smart-seq2. Importantly, it requires no changes to existing Smart-seq2 pipelines, except for the addition of targeting primers during cDNA amplification.

### Simultaneous mapping of mitochondrial and genomic mutations permits high-confidence tracking of leukemic, pre-leukemic and healthy clones

To investigate if MutaSeq can distinguish leukemic, pre-leukemic and residual healthy clones, we generated data from four AML patients with heterogeneous genotypes and phenotypes (Figure 2a, and see also Figures 3b, S2). To allow for a better characterization of stem cells in each patient, cells were sorted such that putative stem and progenitor cells (CD34+) and putatively more mature cells (CD34-) were approximately covered at equal portions (Figure S2). Of note, two of the patients (P2, P4), exhibited bone marrow consisting of >99% of CD34-cells (Figure S2a). Due to different capture rates across individuals and gates, the final data set consisted of between 618 to 1430 cells per patient, of which between 190 and 968 were CD34+ (Figure S2b,c). We therefore avoid statements on quantitative shifts in population size between patients throughout this manuscript.

**Figure 2.**
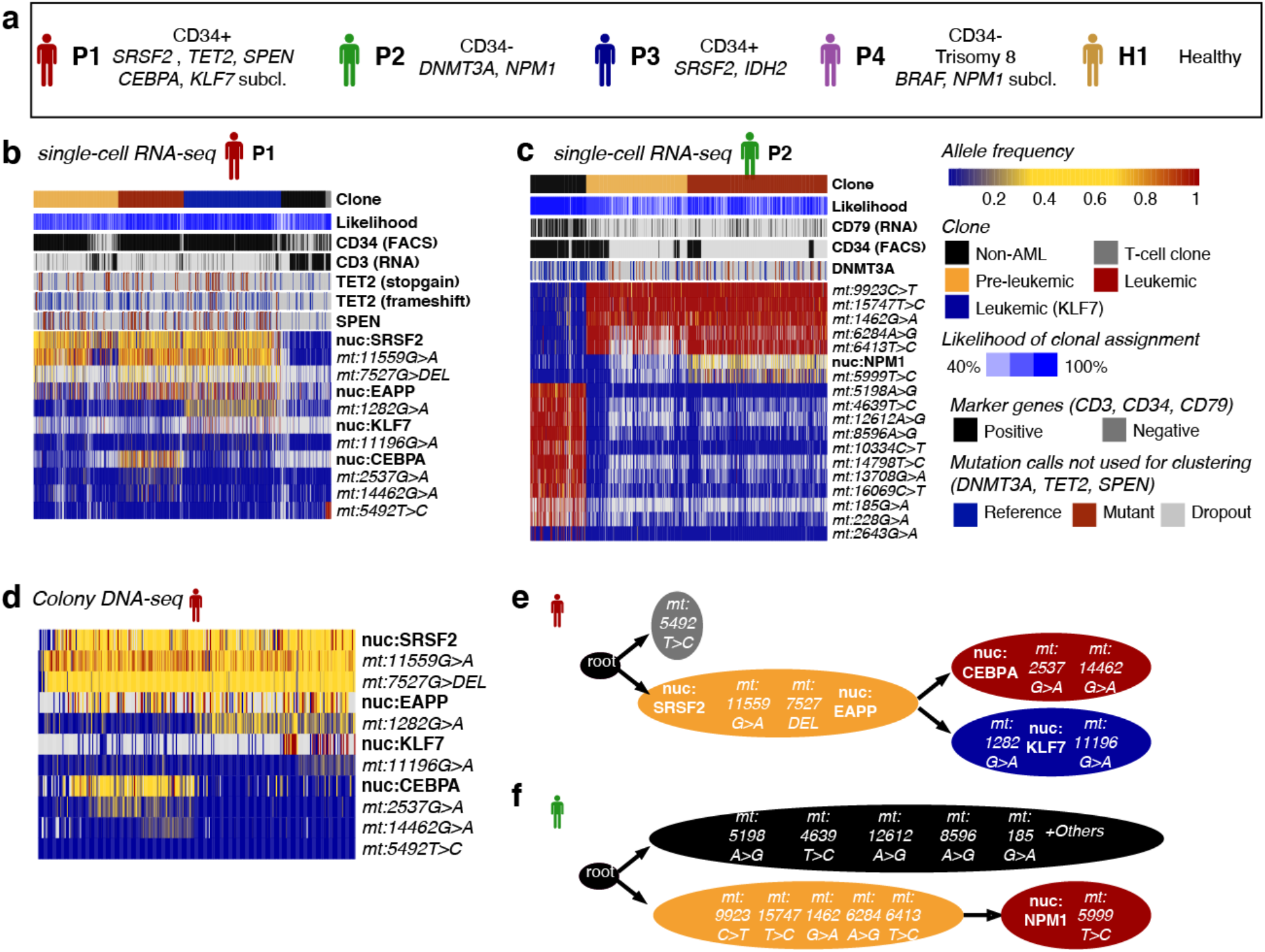
Mitochondrial mutations serve as high-confidence clonal markers in AML. See also Figures S2-S6. a. Overview over the patients. CD34+ and CD34-indicate the dominant surface phenotype of the leukemic blasts, see Figures S2 and 3b for quantification. For each patient, genes containing mutations are printed in italic, see Table S1 for a complete list. Subcl., sub-clonal mutation. b. Heatmap depicting variant allele frequencies (color coded, see right of Figure 2c for a key) observed in single-cell RNA-seq data of n=1430 cells from P1. Grey indicates missing values. Cells and mutations are arranged according to the clustering result obtained by PhISCS^36^ as described in the Methods, section *Analysis of mitochondrial mutations and reconstruction of clonal hierarchies*. Calculation of the likelihood is described in the same section. Mutations with low coverage (in *DNMT3A*, *TET2*, and *SPEN*) were not included in the clustering and are depicted in the heatmaps as metadata, however, in all cases except TET2 frameshift there is quantitative evidence for their association with specific clones (see figure S5b). For reproducing the computations, see the vignettes accompanying the *mitoClone* package. For mutations, nuc is nuclear genome; mt is mitochondrial genome. c. Like panel b, but using n=1066 cells from P2. d. Heatmap depicting variant allele frequencies (color coded, see legend to the right of Figure 2c) observed in targeted DNA-seq data from n=288 single-cell derived colonies from P1. e. PhISCS^36^ was run on the mutational data from P1 to reconstruct an clonal hierarchy, see Methods section “Analysis of mitochondrial mutations and reconstruction of clonal hierarchies”. Take note that while the order of mutations is based on the PhISCS model, the grouping of mutations into clones is based on an arbitrary cutoff to provide a useful clustering for further analyses. See “Clone” in Figure 2c legend for color codes. f. Like e, but for P2.

**Figure 3.**
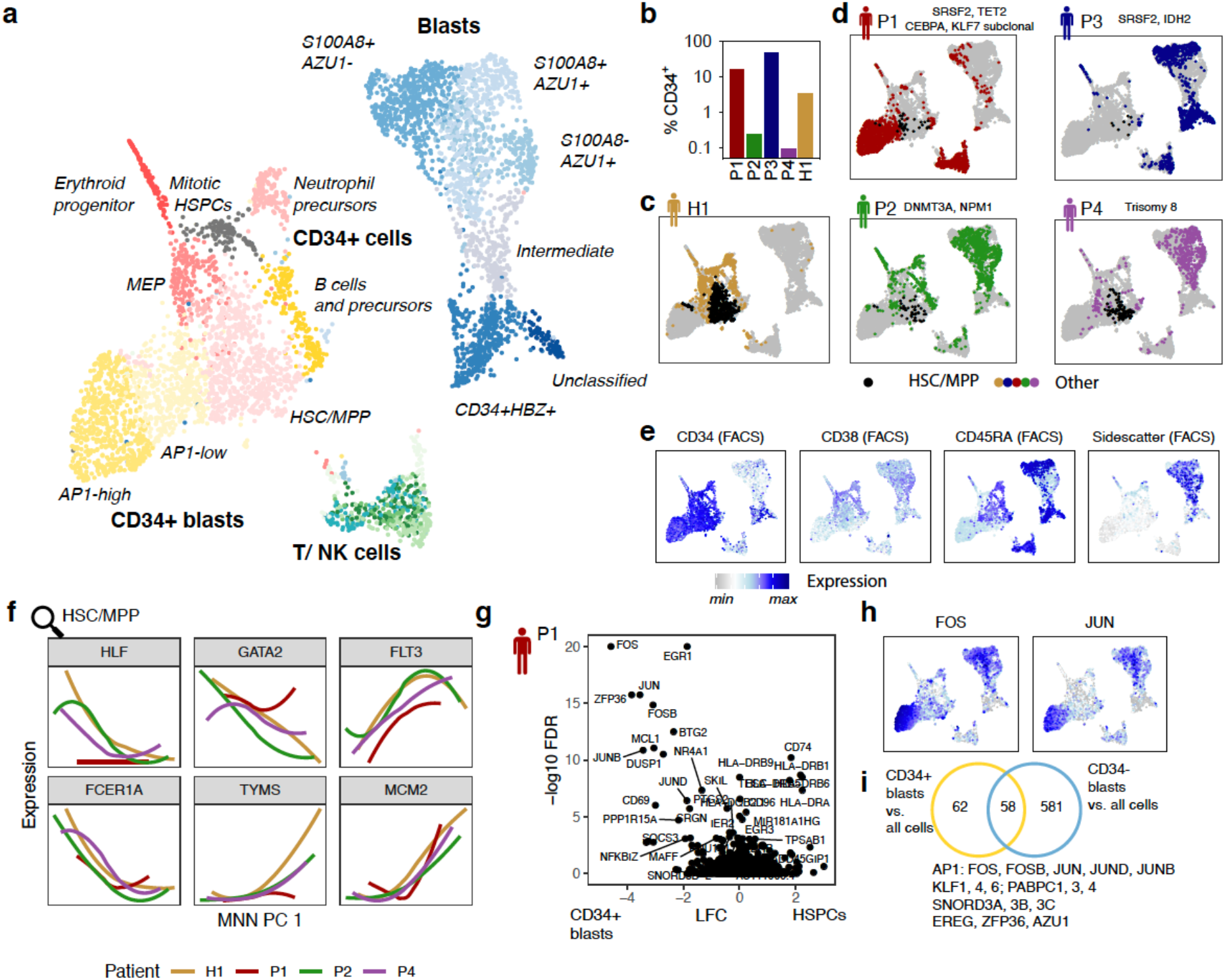
An overview of cell types observed in bone marrow of healthy and leukemic individuals. See also Figures S2 and S7. a. Data from the five individuals (Figure 1a) were integrated using scanorama^59^ and visualized in two dimensions using uMAP^61^. Clusters are color-coded. N = 5228 cells. b. Fraction of CD34+ cells in total bone marrow from the five individuals. CD34+ cells were enriched during FACS sorting, see Figure S2. c. CD34+ cells from a healthy individual^22^ highlighted on the uMAP. Black dots correspond to cells from the HSC/MPP cluster. d. Cells from each patient were highlighted separately on the uMAP. Black dots correspond to cells from the HSC/MPP cluster. e. Logicle-transformed expression of key FACS markers highlighted on the uMAP, see also Figure S7a. f. Data from n= 667 cells from the HSC/MPP cluster were integrated across individuals using MNN^62^. The smoothened expression of several marker genes of healthy HSC/MPP subsets^22^ is plotted over the first dimension of variability identified by MNN. See also Figure S7f. g. Volcano plot of the log10 expression change in n = 569 AP1-high CD34+ blasts vs. n = 667 HSC/MPP-like cells, plotted against corrected p-values from MAST, using a model that accounts for differences in library quality and patient identity / batch (see Methods, section “Single-cell gene expression data analysis” for detail). AP1-high CD34+ blasts were chosen for this comparison since AP1-low blasts, in terms of all marker genes, appear to constitute an intermediate state between Healthy-like HSC/MPPs and AP1-high blasts. h. Log-normalized expression of FOS and JUN on the uMAP from panel a. See panel e for a color scale. i. Venn diagram displaying genes with significant overexpression in AP1-high CD34+ blasts and CD34-blasts, compared to all other cells from the dataset.

We then called nuclear genomic mutations, as well as mitochondrial mutations, at the single-cell level in order to cluster cells into clonal hierarchies. Bulk exome sequencing of the patients had identified known pre-leukemic mutations present at high allele frequency and known leukemic mutations present at a somewhat lower allele frequency (Figure 2a, Table S1. Patient 1: SRSF2,TET2/CEBPA and SRSF2,TET2/KLF7; Patient 2: DNMT3A/NPM1; Patient 3: SRSF2,IDH2; Patient 4: leukemic Trisomy 8 and BRAF mutations). While some statements on clonal hierarchies could be drawn solely based on calls of these nuclear somatic mutation (Figure S3a), the relatively high dropout of these sites impeded robust assignments of cells to clones (Figure S3b-d). Moreover, the result was biased by the expression levels of the mutated genes of interest: in cells with low expression, dropout was higher, leading to a higher fraction of false negative calls, i.e. false classifications of mutant cells as reference (Figure S3c-d and see also Figure 1e). Similar issues were faced by other methods using related approaches of mutation amplification from cDNA^27,28^.

To overcome these limitations, we next determined if mitochondrial mutations can be used to refine clonal hierarchies jointly with the nuclear mutations. Since mitochondrial RNA is extensively edited, it is important to cluster cells based solely on mutations, and not on seemingly polymorphic sites that are the result of post-transcriptional events with no relationship to clonal structure. Here we developed and tested a filtering strategy that only makes use of mitochondrial mutations for clonal tracking if they uniquely occur in individual patients (Figure S4a, see also Methods). This idea assumes that RNA editing events are typically shared between individuals whereas somatic mutations are not. Using the whole exome sequencing data, we validated that this approach, at the level of genomic sites, correctly distinguishes mutations and RNA editing events with a precision of 97% (Figure S4b,c). We further analyzed various control datasets with known associations between mitochondrial mutations and clones^33^ to validate that this approach enables the detection of relevant mitochondrial mutations without a need for a DNA-based reference, and further enables the unsupervised identification of clones (Figure S4d-f). We have implemented all the required filtering and blacklisting routines required for the identification of high-confidence somatic mitochondrial variants in the *mitoClone* R package (https://github.com/veltenlab/mitoClone).

We then computed clonal hierarchies from both nuclear and mitochondrial somatic variants using a mathematical model that accounts for allelic dropout^36^ (see Methods for detail, all required tools are contained in the *mitoClone* package). In patients P1 and P2, pre-leukemic as well as sub-clonal leukemic mutations were significantly associated with distinct sets of well-covered mitochondrial variants (Figure S5a,b), such that clonal hierarchies could be delineated, and a confident assignment of cells to clones became possible (Figure 2b,c and see Figure S3b for a quantitative analysis). Unlike genomic mutation calling from cDNA, identification of clonal identities from mitochondrial mutations is mostly not, and in one case weakly, affected by gene expression levels or library quality (Figure S5c and see also S3c,d), and is possible at lower sequencing depths (Figure S5d-f), since mitochondrial genes are consistently highly expressed.

Importantly, the clonal structure was validated by targeted DNA-sequencing from single-cell derived colonies (Patient 1, Figure 2d). Across the patients, we identified clones carrying known pre-leukemic mutations (e.g. SRSF2, DNMT3A) and sub-clones carrying known leukemic mutations (e.g. CEBPA, NPM1) (Figure 2e,f). Below we functionally characterize these clones as leukemic or pre-leukemic based on their contribution to healthy blood production (see Figure 4).

**Figure 4:**
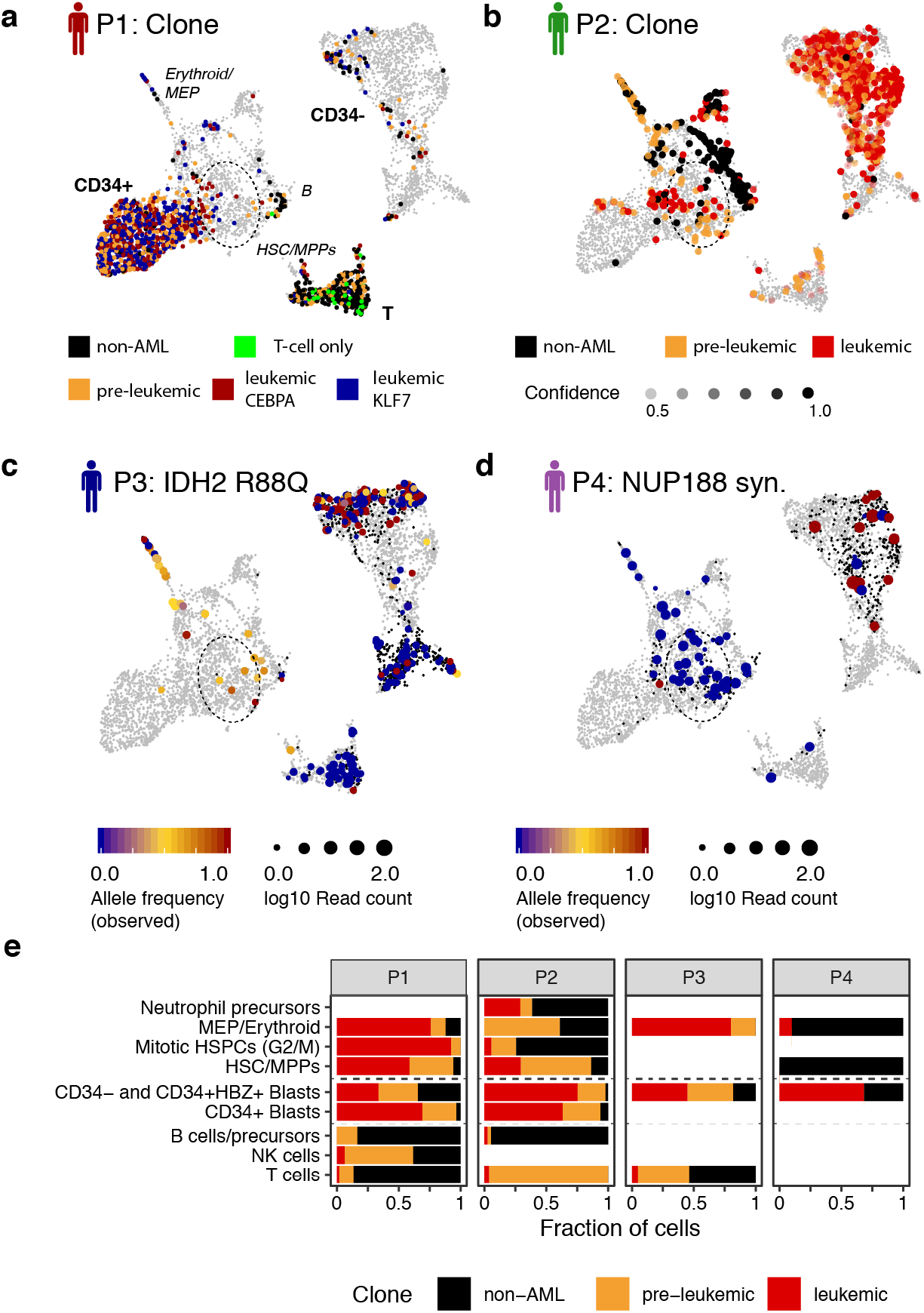
Effects of (pre-)leukemic mutations on cellular differentiation. See also Figure S8. a. Clonal identity of the cells from P1 highlighted on the uMAP; see also Figure 3a and Figure 2b,e. Grey dots correspond to cells from other patients. The dotted ellipse serves as a guide to identify the location of the HSC/MPP population. b. Clonal identity of the cells from P2 highlighted on the uMAP; see also Figure 3a and Figure 2c,f. c. Observed variant allele frequencies for the *IDH2* R88Q mutation from P3 highlighted on the uMAP. Small black dots correspond to cells with no coverage of the mutation. See Figure S8a for an estimate of target capture rates. d. Observed variant allele frequencies for the synonymous *NUP188* mutation from P4 highlighted on the uMAP. This mutation was observed at an allele frequency of 50% in exome data of total bone marrow. Note that CD34+ cells were enriched more than 100-fold during sorting for single-cell RNA-seq, see Figure S2. See Figure S8a for an estimate of target capture rates. e. Estimate of the contribution of different clones to the cell types. For P1 and P2, clonal identities from Figure 2 were used. For P3, cell were classified as leukemic if the *IDH2* mutation was observed, as pre-leukemic if the *SRSF2* mutation was observed, or as non-leukemic if the reference allele was observed for both mutations. Cells without coverage were excluded from the analysis. For P4, cells were classified as leukemic if the *NUP188* mutation was observed, or as non-leukemic if the reference allele was observed. Bars are only shown for populations covered with at least 10 cells. See Figure S8b for absolute numbers and S8c,d for a quantitative analysis.

In patient P3, allele frequencies of leukemic and pre-leukemic mutations were near 50%, indicating the presence of a single leukemic clone (Table S1). In patient P4, an 18-year old individual with a leukemia driven by triplication of chromosome 8, no mitochondrial markers were identified, even though exome sequencing suggested the presence of sub-clonal variants (Table S1). In this case, MutaSeq still permits a qualitative analysis using genomic mutations alone (see below). Taken together, our approach allows for the identification of putatively leukemic, pre-leukemic, and healthy clones and can assign cells to clones with high confidence if mitochondrial somatic variation is present.

### *Identification and characterization of clones* de novo

We next investigated if the use of mitochondrial somatic variants enables the identification and characterization of clones without prior knowledge of nuclear mutations. To that end, we made use of a dataset from patient P1 generated without amplification of nuclear sites (i.e. standard Smart-seq2). A clear clonal structure was identified in an unsupervised manner based solely on mitochondrial variants (Figure S6a). In order to examine whether the presence of somatic genetic variability is associated with the different clones, we then queried the mutational status of 13,797 genomic sites annotated as mutated in AML in the COSMIC database^37^ using a beta-binomial model (see Methods). This unsupervised analysis revealed a highly significant association of the SRSF2 P95H mutation with the leukemic and pre-leukemic sub-clones (Figure S6b,c). Their malignant nature was further evidenced by a markedly reduced ability to contribute to the T cell lineage (Figure S6d and see also below).

To further demonstrate our ability to identify clones *de novo*, we highlight a clonal expansion of non-leukemic cells in P2 (Figure 2f). This clones would have been missed by approaches relying on genomic mutations alone^27,28,31^. Interestingly, these cells were not associated with the pre-leukemic *DNMT3A* mutation. By again querying sites from the COSMIC database^37^ using a beta-binomial model, we identified that they had uniquely acquired a mutation in the *RPL3* gene (Figure S6b,e). These results suggest that this clonal expansion event is independent of the leukemia and associated with the acquisition of unrelated nuclear mutations.

We also take note of a putative non-leukemic clone in P1 marked by a single mitochondrial variant (5492T>C). With one exception, all cells carrying this variant are positive for the T cell marker CD3 (Figure 2b, see also Figure 4a), but they display diverse TCR alpha and beta chain sequences (not shown). Hence, this variant was likely acquired in a T cell precursor, although we cannot formally exclude that it corresponds to a T cell-specific RNA editing event,

Taken together, these results demonstrate that our approach can identify and characterize clones *de novo* without prior knowledge of nuclear genomic mutations. The *mitoClone* package implements all routines for clonal clustering and mutation calling.

### A map of cell states in leukemic bone marrow identifies HSC-like cells

We next used single-cell transcriptome data to distinguish stem cells, progenitor cells and leukemic blasts. To define cell populations in our samples, we integrated the gene expression data from all four patients along with data from CD34+ bone marrow cells of a healthy individual^22^ into a two-dimensional representation, and characterized cell types based on marker gene expression (Figure 3a, S7a,b,c,f, Table S2).

All four bone marrow samples contained cells that clustered with HSCs and multipotent progenitor cells (HSC/MPP) from the healthy individual (Figure 3b-d) and displayed a CD34^+^CD38^low^, FSC/SSC^low^ phenotype (Figure 3e). Unsupervised analysis separated these cells into quiescent immature HSC-like cells (*HLF*+), more proliferative erythromyeloid primed progenitors (*GATA2*+), and lymphomyeloid primed progenitors (*FLT3*+) (Figure 3f, S7f). Cells resembling healthy erythroid progenitors and MEPs were also identified, alongside various types of B-cell precursors and T/NK cells (Figure S7b,c).

In contrast to non-leukemic populations, the majority of cells from leukemic bone marrow samples (‘blasts’) were very different between the patients, in line with previous observations^27^ (Figure 3d): we observed a) differentiated blasts (CD34-SSC/FSC^hi^) that expressed neutrophil genes such as calprotectin (S100A8, S100A9) and *AZU1 (*Figures 3e, S7a); b) CD34+ blasts that were highly mitotic and expressed the fetal hemoglobin *HBZ*; and c) CD34^+^CD38^low^ blasts expressing markers typical of hematopoietic stem and progenitor cells (HSPCs) such as *PROM1* (CD133) and *MEIS1* (Figure S7a,d,e). The latter population appeared to be connected to the HSC/MPP population across a continuum of states that gradually upregulate AP1 transcription factors (FOS, JUN, FOSB, JUNB, JUND) while down-regulating MHC class II (Figure 3g,h, Table S3). A global analysis of highly expressed genes across all populations revealed that high expression levels of AP1, as well as several Krüppel-like factors and poly-A binding proteins, were common to all different blast populations from the patients and distinguished them from healthy progenitors (Figure 3h,i). Of note, high AP1 and KLF activity have recently been identified as a hallmark conserved across genetically distinct types of AML by bulk-sequencing studies that only investigated blast populations^38^. Our results demonstrate that indeed these transcription factors appear to be relevant in all, phenotypically very different, blast populations.

Taken together, all four leukemia samples could be stratified into stem cells, progenitors, and blasts. Furthermore, all patients retain cells highly similar to healthy HSCs, although in variable abundance.

### Clonal tracking identifies cellular differentiation states and gene expression patterns associated with pre-leukemic and leukemic mutations

Gene expression information alone was insufficient to distinguish cancerous from non-cancerous HSPCs (Figure S7f). To definitively characterize cells as (pre-)LSCs or residual healthy cells, we therefore integrated single-cell gene expression data and clonal tracking results (Figure 4a-d). If mitochondrial somatic variability was present, we were able to assign clonal identities with high confidence, allowing us to draw quantitative statements (Patients P1+P2). In the absence of mitochondrial somatic variability, we used nuclear mutation calls in *SRSF2*, *IDH2* and *NUP188* for purely qualitative statements (Patients P3+P4). The capture rates of these marker sites ranged from 70% (SRSF2) to 11% (NUP188) (Figure S8a).

Clones associated with leukemic mutations were most prevalent in the blast compartments and were also detected in the HSPC compartment, but were almost absent in lymphoid (B, NK, T) lineages. In contrast, clones associated with pre-leukemic mutations were found in all lineages, but mostly displayed a decreased prevalence in lymphoid lineages (Figure 4e, S8b-d). These observations confirm the designation of these clones as ‘leukemic’ and ‘pre-leukemic’. Furthermore, these results highlight that the leukemic mutations may initiate differentiation blocks at various levels, as previously reported ^4,27,39^. For example, in patients P1 and P3, leukemic cells had retained the ability to contribute to the erythroid lineage, while in patient P2, this activity was restricted to the pre-leukemic and non-leukemic clones. Importantly, the leukemic cells in P1, P2 and P3 were observed in a cell state that is highly reminiscent of healthy HSCs/MPPs on a molecular level, and retains the ability to contribute to various lineages, i.e. is functionally multipotent.

Next, we investigated the molecular effects of distinct mutations in detail. Previously, the consequences of specific leukemic mutations were commonly studied in mouse models. MutaSeq data allows us to compare gene expression between clones differing only in a single mutation, thereby elucidating the specific effects of that mutation on human hematopoiesis.

Mutations in the *de novo* DNA methyl transferase *DNMT3A* are the most common cause of benign clonal expansions of HSCs in individuals of advanced age^6,7^. In patient P2, *DNMT3A*-mutated pre-leukemic HSPCs were rather stem-like (with relatively high expression of *HLF*) or primed into the erythromyeloid direction (with relatively high expression of *GATA2*) (Figure 4b, 5a). These results are in line with recent findings from DNMT3A knock out mouse models^40^. Interestingly, independent of cell state, this coincided with an upregulation of *MLLT3*, a gene whose enforced expression promotes erythroid-megakaryocytic output from HSCs ^41^ (Figure 5b,c, Table S3).

**Figure 5:**
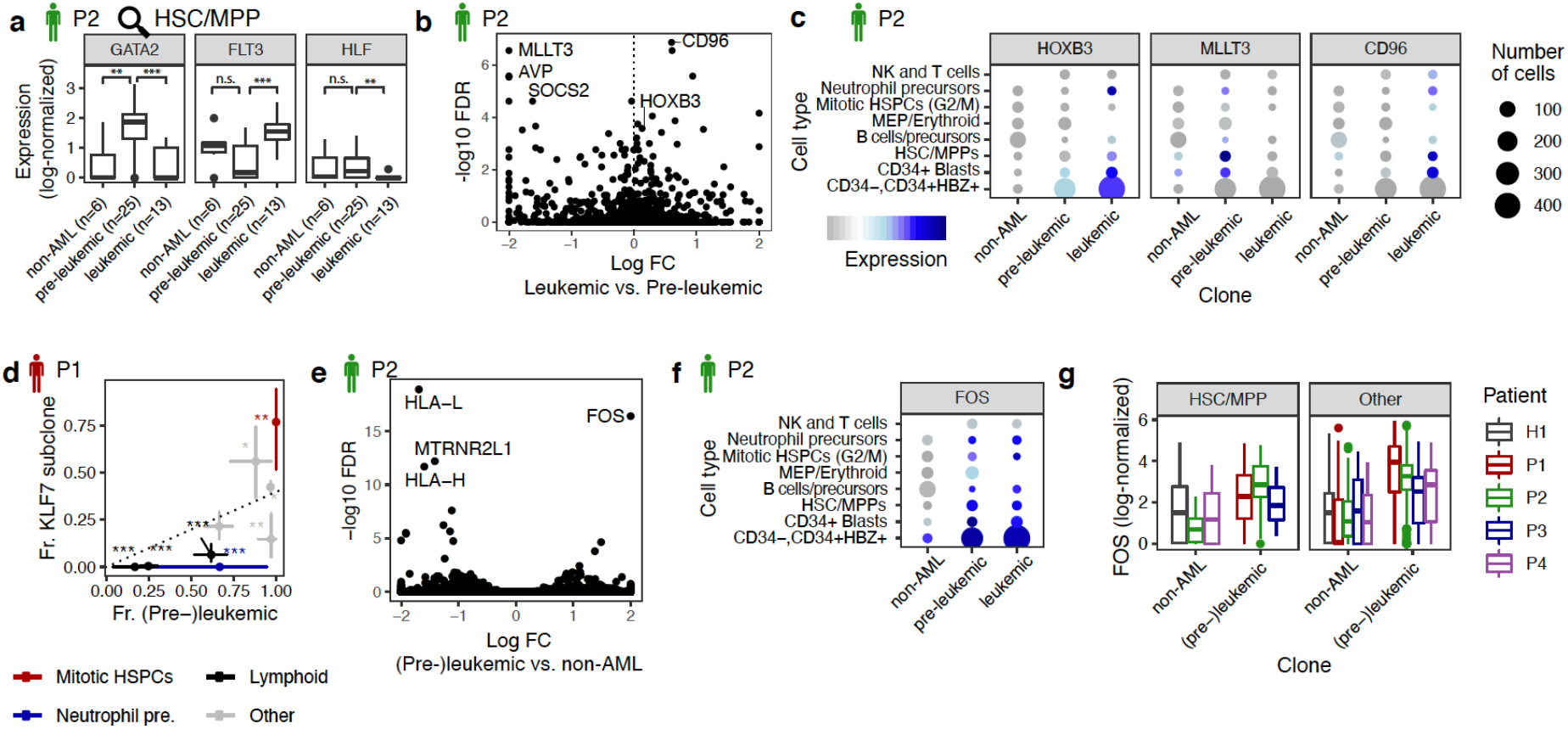
Molecular consequences of leukemic and pre-leukemic mutations. See also Figure S8. a. Log-normalized expression of GATA2, FLT3 and HLF in HSC/MPPs from P2, stratified by clonal identity. See methods, section *Single-cell gene expression data analysis* for detail on data normalization. Asterisk indicate significance from a Wilcoxon test, ***: p < 0.001, **: p<0.01, n.s.: not significant. b. Volcano plot of the log10 expression change in pre-leukemic (n=55) vs. leukemic (n=50) CD34+ cells of P2, plotted against p-values from MAST, using a model that accounts for differences in cell type and library quality (see Methods for detail). Only the following CD34+ cell types from Figure 3a were included in the test: HSC/MPP, CD34+ Blasts (both subsets), Neutrophil precursors, and MEPs. c. Dot plot comparing the expression of relevant genes across non-leukemic, pre-leukemic and leukemic cells in the different cell types. Symbol size scales with the number of cells per cell type. d. Fraction of (pre-)leukemic in relation to the fraction of cells from the KLF7 clone in various cell types in P1. Dotted line indicates the mean ratio across all cells, error bars denote 95% confidence intervals from a beta distribution, and asterisk indicate significant deviation from the mean ratio, as follows: *: p < 0.05; **: p < 0.01; ***: p < 0.001. p-values were computed from the quantiles of a beta distribution against the null hypothesis that the true ratio is the mean ratio observed across all cells. See also Figure S8c. e. Volcano plot of the log10 expression change in (pre-)leukemic (n=105) vs. non-leukemic (n=41) CD34+ cells of P3, plotted against p-values from MAST, using a model that accounts for differences in cell type and library quality (see Methods for detail). Only the following cell types from Figure 3a were included in the test: HSC/MPP-like, CD34+ Blasts (both subsets), Neutrophil precursors, and MEPs. f. Dot plot comparing the expression of FOS across non-leukemic, pre-leukemic and leukemic cells in the different cell types. Dot size and color represent the quantity of cells and gene expression level, respectively (see legend in Figure 5c) g. Boxplots comparing the log-normalized expression levels of FOS between cells with evidence of originating from the non-leukemic clone(s), and cells with evidence of originating from the leukemic or pre-leukemic clones. Cells were assigned to clones as in Figure 4e. See methods, section *Single-cell gene expression data analysis* for detail on data normalization.

Mutations in the multifunctional ribonucleoprotein *NPM1* are identified as drivers for acute myeloid leukemia in 30% of patients and frequently co-occur with pre-leukemic *DNMT3A* mutations^21^. In patient P2, the erythromyeloid bias of the *DNMT3A* clone was lost upon acquiring the leukemic *NPM1* mutation. *NPM1* mutated cells upregulated *HOXB3,* again in line with data from mouse models^42–44^ (Figure 5b,c). Independent of cell state, these cells further exhibited upregulation of *CD96*, which has previously been identified as a leukemia stem cell specific marker^12^. *CD96* was also highly expressed on leukemic HSC/MPP-like cells of patient P3, but not in patient P1, further illustrating the patient-specific nature of LSC markers (Figure S8e).

Mutations in *KLF7* are not commonly observed in leukemia; however, krüppel-like factors are highly expressed by genetically and phenotypically different blasts (see above and ref. ^38^). In patient P1, the KLF7 mutated clone displayed a higher proportion of cells in G2/M phase (Figure 5d). Based on a reanalysis of ATAC-seq data from human CD34+ cells^45^ we found that enhancers containing KLF7 binding sites were enriched near tumor suppressor genes^46^ including *CDK6*, *PTEN*, *RUNX1* and *FLT3*, compared to active enhancers not containing KLF7 binding sites (p = 0.002, Figure S8f).

In sum, we have used a small heterogeneous patient cohort to demonstrate, as a proof-of-concept, that the acquisition of specific mutations is frequently linked to an altered gene expression program, which is consistent with data obtained from mouse models^40,42–44^.

Finally, we compared gene expression between all (pre-)leukemic and non-leukemic cells, with the goal of identifying potential markers or drug targets present in all (pre-)leukemic cells, but not in residual healthy cells. In patient P2, our analysis identified a small number of hits (Figure 5e, Table S3), most notably *FOS*, which was expressed in all (pre-)leukemic cells across cell types, but not by non-leukemic HSC/MPPs (Figure 5f), and the MHC class I genes *HLA-L and HLA-H*, which were expressed by all healthy cells, but not by (pre-)leukemic cells. Across all patients, *FOS* was consistently overexpressed in cells carrying (pre-)leukemic lesions, both in HSCs/MPPs, and in other cell types (Figure 5g).

Taken together, these results demonstrate the ability of MutaSeq and *mitoClone* to delineate developmental and molecular effects of clonal evolution caused by leukemic and pre-leukemic mutations.

## Discussion

Herein, we have described a joint single-cell transcriptomics and clonal tracking approach (MutaSeq and *mitoClone*) for characterizing LSCs, charting their differentiation capabilities, and mapping the molecular consequences of oncogenic mutations. While single-cell gene expression profiling permits the identification of cells with a stem-cell signature, clonal tracking using genomic and mitochondrial mutations allows for a clean separation between healthy and cancerous clones. Thereby we distinguish LSCs, HSCs, pre-LSCs, healthy progenitors, and blasts. We have demonstrated this approach in the context of acute myeloid leukemia, and we propose that similar approaches may be applied to other types of cancers.

By applying our approach to bone marrow samples from four AML patients, we have demonstrated its capabilities:

### Charting the differentiation capacities of hematopoietic clones

By quantifying the contribution of (pre-)leukemic cells to blood lineages, we have shown that in patients P1-P3, leukemic clones not only form blasts but also exist in an HSC-like state. Pre-leukemic clones additionally contribute to erythroid and lymphoid lineages. Multipotent HSCs are therefore the likely cell of origin in these patients^4^. In patient P4, with one exception, only cells with a mature phenotype displayed leukemic mutations, illustrating that the disease can also be fueled by cells with a committed phenotype^39^. Alternatively, the LSC population in this patient might be rare among CD34+ cells.

### Identification of clones de novo

In the presence of mitochondrial somatic variability, our approach does not rely on previously known nuclear mutations to detect clones. In the example of patient 2, we have thereby identified an expanded clone present at very low frequency in total bone marrow, but highly abundant in the CD34+ fraction. We have demonstrated that this clone is associated with a nuclear mutation in the *RPL3* gene, which we discovered *de novo*. This result illustrates the low clonal complexity of hematopoiesis in individuals of advanced age, which often is not associated with candidate driver mutations^20^.

### Identification and characterization of LSCs

Importantly, our approach was capable of identifying leukemic cells highly reminiscent of healthy HSCs. These cells will be important to target therapeutically without ablating their healthy counterparts^9–11^. While stemming from a study cohort of limited size, our data suggest that *FOS* might constitute a potential target for further investigation, as it is expressed throughout the disparate leukemic populations including the most HSC-like cells, pre-LSCs, and blasts, but not in healthy HSCs. Further, we have confirmed that CD96 is a specific marker for LSCs in some patients^12^. Studies in larger cohorts will be required to assess how generally applicable these findings are.

Taken together, our study expands upon earlier work on the molecular phenotypes of AML blasts^27,38^ and constitutes the first detailed characterization of LSCs by single-cell transcriptomics. This advance is owed to three crucial aspects of experimental design.

### Enrichment of relevant starting populations

LSCs are excessively rare and generally present at below 0.1% of total bone marrow^16^. In order to characterize these cells by single-cell genomics, a prior enrichment, e.g. by sorting for CD34 expression, is essential. In future studies, this approach can be complemented with markers that also label CD34-LSCs, such as GRP56^18^.

### Deep transcriptome sequencing

In some cases, the bulk of leukemic cells displays gene expression signatures highly similar to stem cells, as observed here for the CD34+ blasts of patient P1. The differences between LSCs and residual healthy HSCs are even more subtle. Previous work using shallow, microwell based sequencing of AML cells^27^ has not identified differences between LSCs, CD34+ blasts and residual healthy HSCs.

### High-confidence clonal tracking

When available, the use of mitochondrial variants enables the confident assignment of cells to clones, and, thereby, a quantitative analysis of gene expression. In two out of four AML patients, we identified high-confidence, specific mitochondrial genetic markers for pre-leukemic and leukemic sub-clones. In the third patient, allele frequencies of leukemic and pre-leukemic mutations were around 50%, indicating dominance of a single leukemic clone. In the final patient, sub-clonal genomic mutations were observed, but not associated with mitochondrial variability. Interestingly, this patient was only 18 years old and exhibited a leukemia possibly driven by a ‘catastrophic’ triplication of chromosome 8. The length of the pre-leukemic phase, the buildup of mitochondrial variants accumulated during normal ageing, as well as unknown factors affecting mitochondrial mutation rates might all contribute to the presence of mitochondrial marker mutations.

In the context of recent methods for clonal tracking within single cell transcriptomics^27–29,31,33^, the use of the MutaSeq protocol and *mitoClone* computational pipeline combines the strengths of previous approaches relying on either nuclear or mitochondrial variants, but also has limitations. Specifically, droplet- or microwell-based protocols for single-cell RNA-seq and clonal tracking^27–29^ provide low coverage of mitochondrial genomes and high dropout of nuclear genomic sites, and therefore do not enable a confident assignment of cells to clones. Single-cell ATAC-sequencing^33,34,47^ and Smart-seq2 efficiently cover mitochondrial genomes, but their coverage of nuclear genomic sites is absent or low, hardening interpretability of the data. Finally, while TARGET-seq addresses the limitation of dropout of nuclear sites, depending on the implementation it does not offer sufficient mitochondrial coverage, or is of very limited throughput. Of all methods available to date, the approach presented here converges on crucial aspects, specifically: a) high mitochondrial coverage, allowing us to identify benign expanded clones *de novo*, even in the absence of known genetic markers, b) decreased dropout of relevant genomic mutations, permitting the association of clones with genomic mutations, if present and c) a highly confident assignment of cells to clones, enabling quantitative analyses of clone-specific gene expression (Figure S9). These capabilities expand the potential applications of our approach to the study of clonal dynamics during ageing and oncogenesis beyond the hematopoietic system.

### Limitations

The major limitation of our pipeline is that it requires natural somatic variability to resolve clones at high confidence. In the absence of mitochondrial somatic variation, the MutaSeq protocol can be used to draw qualitative statements on clonal differentiation capacities (similar to ref. ^28^, and with an improvement over Smart-seq2), but due to dropout neither enables the statistically confident assignment of cells to clones, nor differential expression testing between clones. In the presence of mitochondrial somatic variation, an association between clones and nuclear mutations is only possible for mutations in highly expressed genes, and can be limited by the nature of the mutation (e.g. frameshift mutations) and possibly other factors such as sequence complexity. The third limitation of MutaSeq is its relatively low throughput. This limitation is currently shared with Smart-seq2 and alternative single-cell RNA-seq methods allowing high-confident assignment of cells to clones^31,33^. Future work will focus on the inclusion of full-length coverage of the mitochondrial genome in droplet-based single-cell RNA-seq platforms.

## Material and Methods

### Patient and sample collection

The AML samples were collected from diagnostic bone marrow aspirations at the University hospitals in Heidelberg, Germany, and Mannheim, Germany after obtaining informed written consent. Bone marrow mononuclear cells were isolated by density gradient centrifugation and stored in liquid nitrogen until further use. All experiments involving human samples were conducted in compliance with the Declaration of Helsinki and all relevant ethical regulations and were approved by the ethics committees of the medical faculties Heidelberg and Heidelberg-Mannheim of the University of Heidelberg.

### Deep exome sequencing and target selection

For exome sequencing, DNA was extracted from 9×10^3^ flow sorted CD34+ cells (for CD34+ leukemias: P1, P3) or total bone marrow. As healthy controls, we used a buccal swap (P1), FACS-sorted CD45-CD105+ MSCs (P3) or *in vitro* expanded MSCs (P2 and P4)^5^. Sequencing libraries were constructed using the SureSelect HS XT Target Enrichment System v6 (Agilent), and a mean on-exon sequencing coverage of at least >70X was obtained for each patient. Genomic alignments were performed using BWA MEM v0.7.15^49^ and cancer variants were identified using Mutect2 v3.8 (P1 and P3) and v4.0.9 (P2 and P4)^50^, following the GATK best practice recommendations. Variants were annotated using ANNOVAR^51^. Output from Mutect2 was filtered to remove variants that did not overlap with known genes. The final list of candidate variants included only those with allele frequencies (AF) greater than 4% in the cancer exome sample and with an AF four-fold larger than in the healthy exome sample (Table S1). Finally, the candidates for targeting were hand-selected from this list with a focus on cancer relevant genes, highly expressed genes, and potential sub-clonal markers.

### FACS sorting

Bone marrow mononuclear cells were stained for 30 minutes on ice according to standard protocols. For single-cell liquid cultures and MutaSeq, cells were stained with fluorescent-labelled antibodies against lineage markers (CD4, CD8, CD19, CD20, CD41a, CD235a) and additional markers (CD45RA, CD135, GPR56, CD34, CD38, CD90, CD33, Tim3), and sorted according to the gating scheme illustrated in Figure S2. BD FACS Fusion (BD Biosciences) equipped with 405nm, 488nm, 561nm and 640nm lasers were used. Of note, in P1 and P4, Lin+ cells could not be efficiently processed into libraries for unknown reasons and are therefore excluded. A list of all antibodies used can be found in Table S4.

### Primer Design

Primers for MutaSeq, for other single-cell targeting protocols tested (Figure S1), as well as for targeted DNA sequencing were designed using the computational pipeline available at http://git.embl.de/velten/PrimerDesign. For MutaSeq, the refgene transcripts spanning each genomic site of interest were selected as template; if multiple refgene transcripts were found for one site, a consensus transcript containing only exonic sequences present in all variants was created. We then used primer3^52^ to design five possible pairs of primers for each intended target, with an amplicon length of 90-145bp and a melting temperature of (nominally) 60°C. BLAST was used to remove primer pairs which potentially form off-target amplicons. Then, the pair complementarity (i.e. potential to form dimers) was computed for each possible combination of primers across all target sites (forward-reverse, forward-forward and reverse-reverse). In initial experiments, primers with high complementarity scores were found to abundantly form primer dimers (not shown). In order to identify a set of primers that covered the maximal number of genes while strictly forbidding primers with high complementarity scores, a graph was constructed that connected all primers with different targets to each other if their complementarity score was lower than 15. A maximum clique-finding algorithm^53^ was then used to identify the largest mutually connected component in the graph. Thereby, the largest number of targets that efficiently avoids dimer formation was selected.

Nextera adapters were added to all primers designed accordingly (fwd: GTCGTCGGCAGCGTCAGATGTGTATAAGAGACAG, rev: GTCTCGTGGGCTCGGAGATGTGTATAAGAGACAG).

For targeted DNA-seq experiments, the genomic sequence surrounding the target was used as template and nested PCR primers were designed. Inner primers were designed as in the case of MutaSeq, and outer primers surrounding the inner PCR product with an amplicon length of 200-350bp and a nominal annealing temperature of 58°C were added. A list of all primers used for this study is included in Table S5.

### Single-cell RNA sequencing with targeting of genomic sites of interest (MutaSeq)

MutaSeq is based on the Smart-seq2 protocol^54,55^ with the modifications introduced by^22^. For lysis, we used 5μL of a buffer containing 0.1μL RNAsin+ (Promega), 0.04μL 10% Triton X-100 (SigmaAldrich), 0.1μL of 100μM Smart-seq2 Oligo-dT primer (SigmaAldrich) and 1μL dNTP mix (10mM each, NEB). In P1, we had additionally included 0.075μL of a 1:1,000,000 dilution of ERCC spike-in mix 1 (Ambion), as well as a control spike-in to quantify the false positive detection rate of mutations (see Figure S10). Plates were snap frozen directly after sorting and later thawed at 10°C in a PCR machine for 5’ and denatured at 72°C for 3’. 5μL of a buffer containing 0.25μL RNAsin+, 2μL 5x SMART FS buffer, 0.5μL DTT 20mM, 1μL SmartScribe enzyme (all TaKaRa) and 0.2μL 50μM Smart-seq2 TSO (Exiqon) were then added and RT was performed for 90’ at 42°C, 10 cycles of [50°C, 2’ and 42°C, 2’], and enzyme inactivation at 70°C for 15’. Then, we added 15μL PCR mix containing 12.5μL KAPA HiFi HS mastermix (Merck), 0.25μL 10μM Smart-seq2 ISPCR primer (SigmaAldrich) and 0.5μL of a pool of all targeting primers, present at 1μM each. cDNA amplification was performed by 98°C 3’, 21 cycles of [98°C, 20’’, 67°C, **60’’**, 72°C 6’] and 72°C, 5’. cDNA was the cleaned up using an equal volume (25μL) of CleanPCR beads (CleanNA) and tagmented using homemade Tn5^56^.

### Single-cell cultures

Bone Marrow mononuclear cells from patient P1 were stained and Lin- or Lin-CD34+ single cells were index-sorted into ultra-low attachment 96-well plates (Corning) containing 100μL StemSpan SFEM media (Stem Cell Technologies). Media was supplemented with penicillin/streptomycin (100ng/mL), L-glutamine (100ng/mL) and the following human cytokines (all from Peprotech): SCF (20ng/mL), Flt3-L (20ng/mL), TPO (50ng/mL), IL-3 (20ng/mL), IL-6 (20ng/mL), G-CSF (20ng/mL), EPO (40ng/mL), IL-5 (20ng/mL), M-CSF (20ng/mL), GM-CSF (50ng/mL). After 21 days at 5% CO_2_ and 37°C, colonies were imaged by microscopy, and processed as detailed in the following.

### Targeted DNA sequencing by nested PCR amplification

Single-cell derived colonies were transferred into 50μL buffer RLT (Qiagen). Cleanup was performed using CleanPCR beads (CleanNA) at a 1.8x volume ratio and eluted in 20μL 10mM Tris-HCl pH 7.8. 4.5μL were transferred to a PCR plate containing 7.5μL Kapa HiFi HS mastermix and 3 μL of a pool of all outer primers (Table S5, each primer at 0.5μM) were added, followed by a PCR program of 98°C 3’, 30 cycles of [98°C, 20’’, 63°C, 60’’, 72°C 10’’] and 72°C, 5’ and subsequent enzymatic cleanup with 2.5μL 10x ExoI buffer, 0.4μL ExoI (NEB) and 0.4μL FastAP (ThermoFisher), 30’ incubation at 37°C and 5’ inactivation at 95°C. Afterwards, 1μL was transferred to a PCR tube containing 5.9μL water, 7.5μL Kapa HiFi HS mastermix and 0.6μL of a pool of all inner primers (Table S5, each primer at 0.5μM), followed by a PCR program of 98°C 3’, 15 cycles of [98°C, 20’’, 65°C, 15’’, 72°C 30’’], 72°C, 5’ and enzymatic cleanup as above. 1μL was then transferred into a PCR with Nextera indexing primers as described in^56^ and amplified with 98°C 3’, 10 cycles of [98°C, 20’’, 60°C, 15’’, 72°C 30’’] and 72°C, 5’.

### Processing of Next Generation Sequencing data

Raw sequencing reads from MutaSeq and Smart-seq2 experiments were processed using the BBDuk software to trim both the standard Illumina Nextera adapters and the ISPCR adapter. Reads were then mapped to the hg38 human genome (Ensembl release 95) using STAR v2.6 (Dobin et al, 2013), with the outFilterMismatchNmax parameter set to 5. Exonic gene counts were tabulated, keeping only reads that did not overlap with targeted regions, overlapped with only one annotated gene, and with lengths greater than 30 nt. For the colony DNA sequencing experiment, reads were mapped to the hg38 human genome using bwa mem (v0.7.17)^49^.

### Analysis of mitochondrial mutations and reconstruction of clonal hierarchies (mitoClone package)

The following set of routines are implemented in the mitoClone package available form https://github.com/veltenlab/mitoClone, and documented further in the package vignettes.

### Construction of allele count tables

For each cell and position of mitochondrial genome as well as other genomic sites of interest, count tables for all nucleotides (A/C/G/T), and deletions were created from the BAM files. To this end, the *mitoClone* package implements the *baseCountsFromBamList* function, which essentially serves as a wrapper to the bam2R function from the deepSNV R package^57^.

### Filtering of mitochondrial variants

To identify relevant somatic variants, we implemented the *mutationCallsFromCohort* function. In short, we select coordinates in the mitochondrial genome containing at least 5 reads each in at least 20 cells. To distinguish RNA editing events and true mitochondrial mutations at the level of genomic sites, we then identify mitochondrial variants that occur in several individuals. For this purpose only, individual cells are therefore called as “mutant” in a given site of the mitochondrial genome if at least 10% of the reads from that cell were from a minor allele (i.e. distinct from the reference). Mutations present in at least 1% of cells in a given patient, but no more than 10 cells in any other individual, are then included into the final dataset and counts supporting the reference and mutant alleles are computed as for sites of interest in the nuclear genome. Mutations present in several individuals are stored as a blacklist and were used further for filtering some of the data analyzed in Figure S4. Importantly, the result from this step is simply a list of genomic sites that are likely to display genetic variability across single cells. The vignette ‘Variant calling and blacklist creation of the *mitoClone* package provides further recommendation for the choice of filtering parameters.

### Construction of clonal hierarchies

To construct a basic clonal hierarchy, we implemented the *muta_cluster* function. In short, all nuclear and mitochondrial variants with coverage in at least 20% of cells are selected. Observed variant allele frequencies (VAF) are then computed for each cell. From these values, we compute a ternary matrix of observed variant calls *N*_*c,g*_; for each cell *c* and genomic site *g*, cells were classified as mutant (1) if the VAF was above 5%, reference (0) if it was below 5%, or dropout (NA) if the site was not covered. We thereby assign a genotype ‘mutant’ or ‘non-mutant’ to single cells at all genomic sites selected in the filtering step, using a less stringent cutoff for calling the site as mutant. Then, PhISICS^36^ is used to reconstruct a clonal tree. Unlike conventional algorithms for the reconstruction of phylogenetic hierarchies, PhISCS is very robust with regard to noise. For the figures presented in the main text, we ran PhISCS assuming a false positive rate (FPR) of 3% and an allelic dropout rate (AD) of 10% across all genes. We additionally estimated the dropout rate on a per-gene level from the number of complete dropouts, and varied the resulting parameter vector using Latin hypercube sampling around the means using a beta distribution with concentration parameter of 10. Across 80 sampling runs, the same tree was consistently obtained except that the order of nodes within clones was swapped (not shown). PhISCS results are therefore robust to variations in the parameters of the statistical technique used. The false positive rate of MutaSeq had been empirically estimated using spike-in controls (Figure S10).

### Clustering of mutations into clones and assignment of cells to clones

PhISCS provides a maximum likelihood phylogenetic tree, enforcing an ordering of mutations. In reality, however, not all intermediate evolutionary steps are represented by cells present in the biological sample (for example, in P1, there are few or no cells displaying the SRSF2 mutation, but not the mt:11559G>A mutation). Hence, the order of the nodes in the maximum likelihood tree is to some extent arbitrary and driven by noise; even if there is some statistical support for a specific order, it may be attractive in practice to merge mutations into ‘clones’ so as to obtain a biologically meaningful, interpretable analysis. We therefore implemented the *clusterMetaclones* function, which employs a likelihood-based approach. The maximum likelihood tree is split into contiguous linear branches (e.g. for P1, nuc:SRSF2, mt:11559G>A, mt:7527DEL and nuc:EAPP constitute one such branch). Within each branch, all nodes are then swapped with each other and the likelihood of the data given the altered structure is calculated using the PhISCS model:

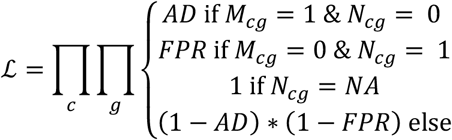

Here, *M*_*c,g*_ indicates whether according to the model, cell *c* is mutant at genomic site *g*.

The branch is then split into clones such that within each clone, the average difference in log-likelihood incurred by swapping nodes was smaller than 1 per cell. This threshold is set arbitrarily to obtain a practically useful grouping of mutations into clones. The vignette ‘Computation of clonal hierarchies and clustering of mutations’ of the mitoClone package provides further practical recommendations. The same model was used to compute the likelihood of clonal assignments for each cell. For the analysis in Figure S3b, this estimate was then transformed to bits of information using the Kullback-Leibler distance:

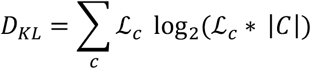

where |*C*| is the total number of clones.

### Quantitative analyses

For all quantitative analyses of clones (differential expression testing, constribution of clones to cell types) cells with a likelihood of clonal assignment of <0.8 were removed. *De novo mutation calling.* To identify nuclear mutations associated with the clones, we performed variant calling at a list of candidate sites from COSMIC^37^. We therefore focused on a subset of COSMIC including putatively pathogenic SNVs and small InDels, variants in expressed genes (mean >20 reads per cell), and variants associated with a primary site “haematopoietic_and_lymphoid_tissue”, obtaining a list of 13,797 sites of interest. SNVs commonly observed as germline variants in the 1000 genomes project dataset were removed^58^. Allele count tables were created for each site as described above. Finally, a beta-binomial model with the same probability for mutant in all cells was compared to a beta-binomial model with a different probabilities for mutant in each clone using Akaike’s Information Criterion.

### Single-cell gene expression data analysis

Cells with less than 500 distinct genes observed and genes that appeared in less than 5 cells were removed. Additionally, data from a healthy individual (“H1”) was downloaded from the NCBI Gene Expression Omnibus (GSE75478). Data from all individuals was then integrated using scanorama according to the workflow described by the authors^59^. The low dimensional data representation obtained by scanorama was then loaded into Seurat^60^, and default Seurat implementations of UMAP^61^ and graph-based clustering were used for data visualization and clustering, respectively, using the first 15 scanorama components.

For more detailed analyses of the T/NK-cell and HSC/MPP populations, cells with these identities were selected and the MNN^62^ data integration workflow was repeated using raw expression counts of these specific cell populations as input.

Differential expression testing was performed using MAST^63^, using a linear model containing the variable of interest (e.g. clonal identity), a library quality covariate (i.e. the number of genes observed per cell), and, when applicable, additional covariates accounting for patient and cell type.

For the display of gene expression values only, data were normalized according to the Seurat defaults (i.e. divided by the total count of RNA in the cell, multiplied by a scale factor of 10,000 and log-transformation).

### Data visualization

All plots were generated using the ggplot2 (v. 3.2.1) and pheatmap (v. 1.0.12) packages in R 3.6.2. Boxplots are defined as follows: the middle line corresponds to the median; lower and upper hinges correspond to first and third quartiles. The upper whisker extends from the hinge to the largest value no further than 1.5 * IQR from the hinge (where IQR is the inter-quartile range, or distance between the first and third quartiles). The lower whisker extends from the hinge to the smallest value at most 1.5 * IQR of the hinge. Data beyond the end of the whiskers are called “outlying” points and are plotted individually^64^.

### Statistics and Reproducibility

Statistical analyses were performed using R. Statistical details for each experiment are provided in the figure legends.

## Data availability

Count tables and other processed data necessary to reproduce all analysis from the manuscript are deposited in figshare with DOI 10.6084/m9.figshare.12382685.v1^65^. Raw sequencing data are deposited under a Data Access Agreement to protect patient privacy in the European Genome-Phenome Archive with the accession id EGAS00001003414. Sequencing data from the healthy individuals are deposited in GEO with the accession id GSE75478.

## Code availability

Code for MutaSeq primer design is available at https://github.com/veltenlab/PrimerDesign. A package for data analysis is available at https://github.com/veltenlab/mitoClone.

## Acknowledgements

We thank Niko Beerenwinkel for discussions, and Andreas Gschwind for contributing code for primer design. We thank the members of the Steinmetz and Haas labs for discussions and support, and we thank the EMBL and DKFZ Genomics core facilities, and the EMBL flow cytometry core facility for technical support. This project was financially supported by the Deutsche José Carreras Leukämie Stiftung grant DJCLS 20 R/2017 (to LV, SH, LMS and AT), the Emerson foundation grant 643577 (to LV and LMS) and the German Bundesministerium für Bildung und Forschung (BMBF) through the Juniorverbund in der Systemmedizin ‘LeukoSyStem’ (FKZ 01ZX1911D to LV, SH and SR). Contributions by SR were further supported by Emmy Noether Fellowship RA 3166/1-1 (DFG). Contributions by CP were supported by a Max-Eder Grant (German Cancer Aid 70111531). Contributions by DN, JCJ, WKH and TB were supported by the Gutermuth Foundation, the H.W. & J. Hector fund, Baden-Württemberg, and the Dr. Rolf M. Schwiete Fund, Mannheim. DN is an endowed professor of the Deutsche José Carreras Leukämie Stiftung (DJCLS H 03/01).

## Author contributions

LV developed MutaSeq and performed single-cell RNA-seq experiments with assistance by JM, DRL and CST. PH performed sample characterizations, single-cell culture experiments and FACS sorting with support by MP, AD, SR and supervision by SH. BAS and LV developed mitoClone and analyzed the data with contributions from RF. LV, SH, AT and LMS conceived the study. LV and BAS wrote the manuscript with contributions from PHM, SH, SR and LMS. All other authors were involved in the collection and initial characterization of samples. All authors have read and commented on the manuscript.

---

**Figure S1.**
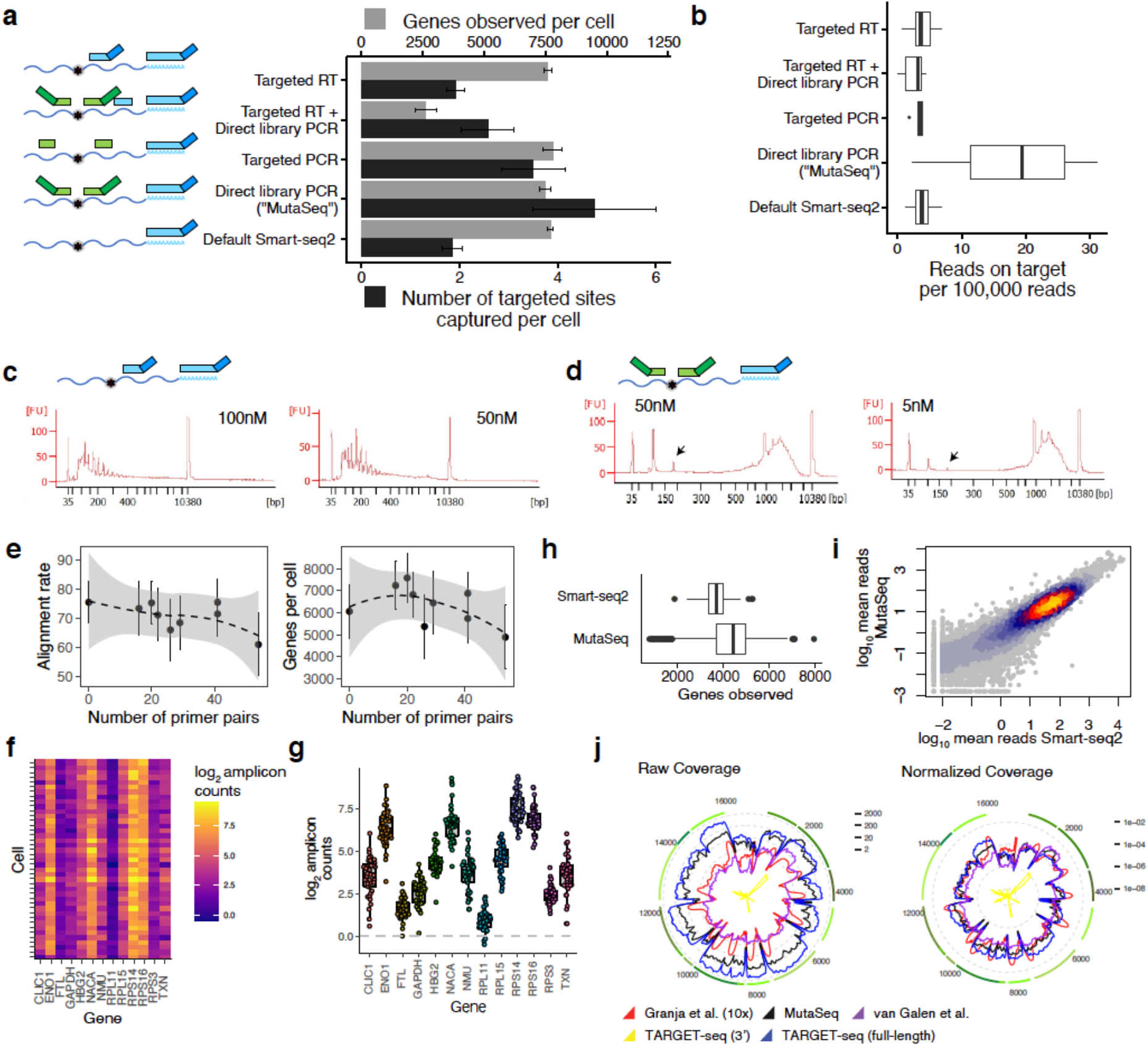
Development of MutaSeq. See also Figure 1. a. Comparison of different strategies for targeting 14 genomic sites of interest during Smart-seq2 library preparation; see Table S5 for primers used. In the ‘targeted RT’ protocol, a reverse transcription primer carrying the ISPCR sequence was placed downstream of the sites of interest. In the ‘targeted RT+ direct library PCR’ protocol, a targeted RT primer without ISPCR sequence was used in conjunction with targeting primers included during PCR. In the ‘targeted PCR’ protocol, PCR primers were used to generate amplicons of 250-350 bases, whereas in the ‘direct library PCR’ protocol, shorter amplicons were used and primers were fused to Nextera sequencing adapters; see also Figure 1b. 8-16 K562 cells were sequenced with each protocol and the number of genes observed per cell as well as the number of target sites covered was quantified. Error bars indicate the standard error of the mean. For all analysis in this panel, reads were down-sampled to 250,000 reads per cell prior to alignment to account for differences in coverage. b. Boxplot comparing the mean number of reads per target in the different protocols. See Methods section on ‘Data Visualization’ for a definition of boxplot elements. c. Bioanalyzer traces for a protocol targeting sites of interest during RT only. Concentrations correspond to the final concentration of targeting primers in the RT reaction. d. Bioanalyzer traces for a protocol targeting sites of interest during library amplification PCR (MutaSeq). Arrows highlight the MutaSeq amplicons. Concentrations correspond to the final concentration of targeting primers in the PCR reaction. e. Primer sets developed for the MutaSeq patients (Table S5) were combined in all possible combinations (i.e. P1+P2 primers, P1+P3+P4 primers, etc.) and libraries were generated from n=8 K562 cells each to evaluate the effect of multiplexing primers on alignment rate (left) and number of genes observed (right). f. Primer pairs were designed surrounding randomly selected sites on 13 highly-expressed genes in K562 cells (Table S5). The MutaSeq protocol was then performed using these primers on n=48 K562 cells. For each gene, the number of reads from MutaSeq amplicons (i.e. complete matches) is shown, after subtracting the average coverage of the surrounding areas outside of the targeted site (i.e. potential background signal). Seven cells with poor alignment rates (below 50%) were removed. g. Amplicon counts for 13 genes across 41 cells is shown as boxplots. The points in the overlaid beeswarm plot represent cells. Same underlying data as used in Figure S1f. h. Number of genes observed per cell, across n=206 (Smart-seq2) or n=658 CD34+ (MutaSeq) cells. See Methods section on ‘Data Visualization’ for a definition of boxplot elements. i. Mean gene expression levels measured by Smart-seq2 are compared to mean gene expression levels measured by MutaSeq. Color reflects point density. j. Logarithmic coverage of the mitochondrial genome compared between different methods^27,31,66^. For the plot on the right, coverage was normalized to the number of reads aligning to the transcriptome.

**Figure S2.**
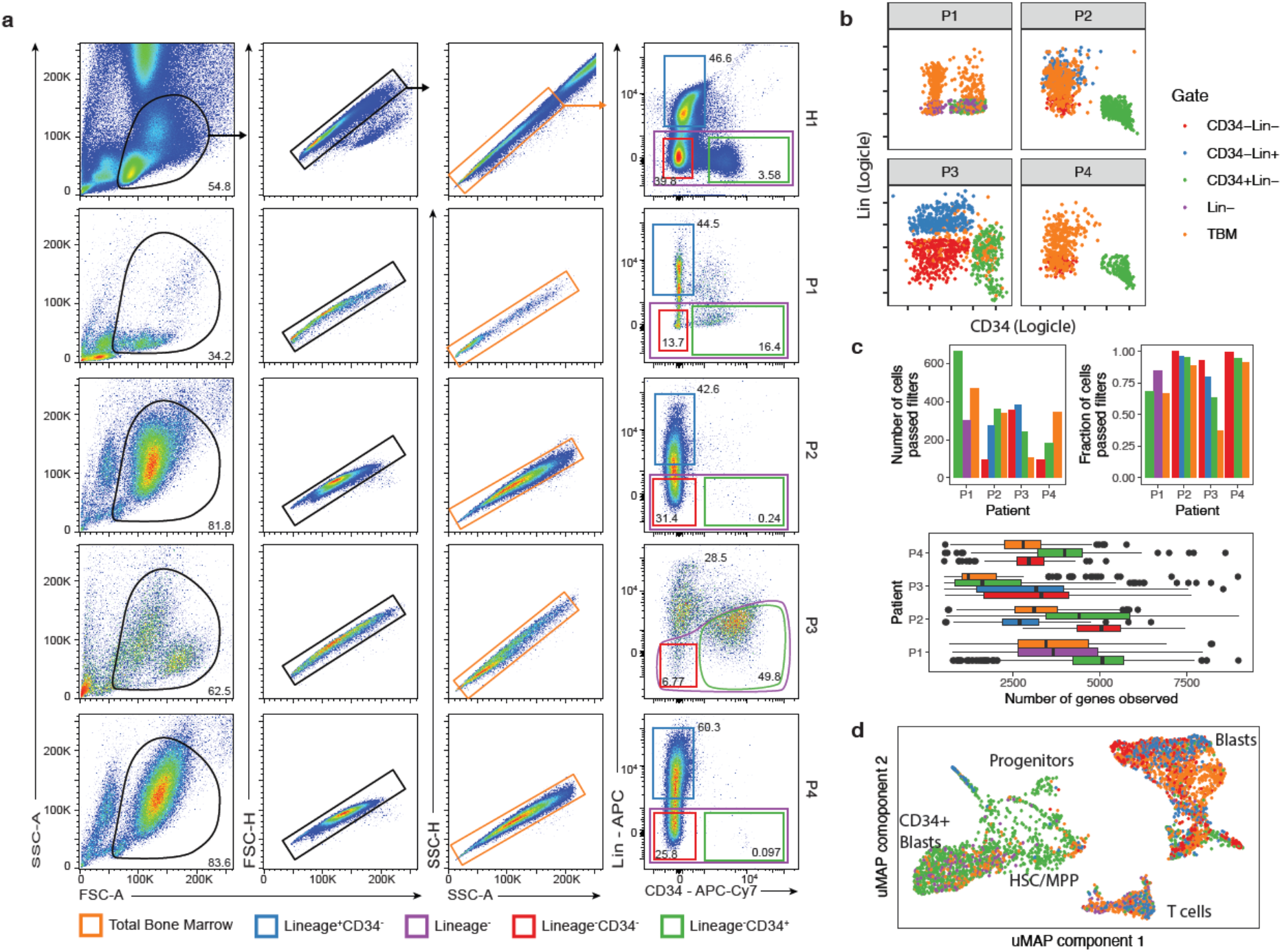
FACS sorting schemes and quality control of single-cell RNA-seq data. See also Figure 2+3. a. Gating schemes used for the different patients. b. Index values of cells included into the final data set c. Top left: number of cells from the various gates included into the final data set; see panel a/b for a color scheme. Top right: fraction of cells passing filters, stratified by patient and gate. Bottom: box plots depicting the number of genes observed in cells from the final data set. Cells with more than 500 genes observed were included. d. uMAP plot of cells from all individuals, with cells color coded according to their sorting gate. See panel a/b for a color scheme, and main Figures 3a-e and S7a for a more detailed description of the uMAP.

**Figure S3.**
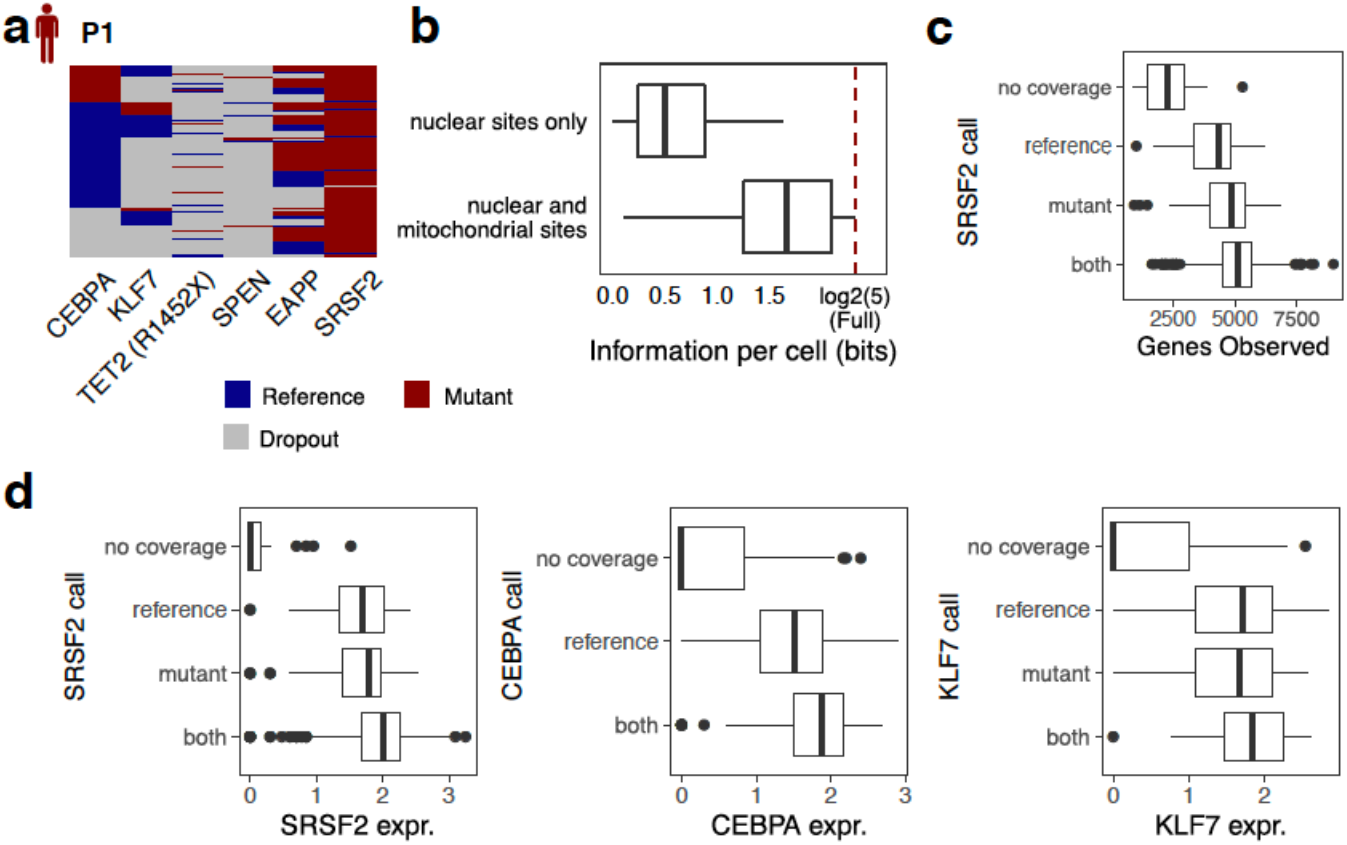
Evaluation of nuclear markers for clonal tracking in single-cell RNA-seq data. See also Figure 2. a. Heatmap depicting mutation calls of nuclear genomic mutations only for P1. Clustering was performed as described in the methods, section *Analysis of mitochondrial mutations and reconstruction of clonal hierarchies*, except that here, only nuclear mutations were included. Rows represent cells. b. The full specification of clonal identities in case of P1 requires log2(5) bits of information, since there are five main clones (Figure 2e). For each cell, the information available from nuclear genomic sites only, or both nuclear and mitochondrial sites, was quantified as described in the Methods, section *Analysis of mitochondrial mutations and reconstruction of clonal hierarchies*. c. Box plots evaluating the extent to which differences in library quality affect measurements of the mutational status of target genes; data from P1 is shown. d. Like panel c, but investigating the effect of the expression of a targeted gene on the ability to measure its mutational status.

**Figure S4.**
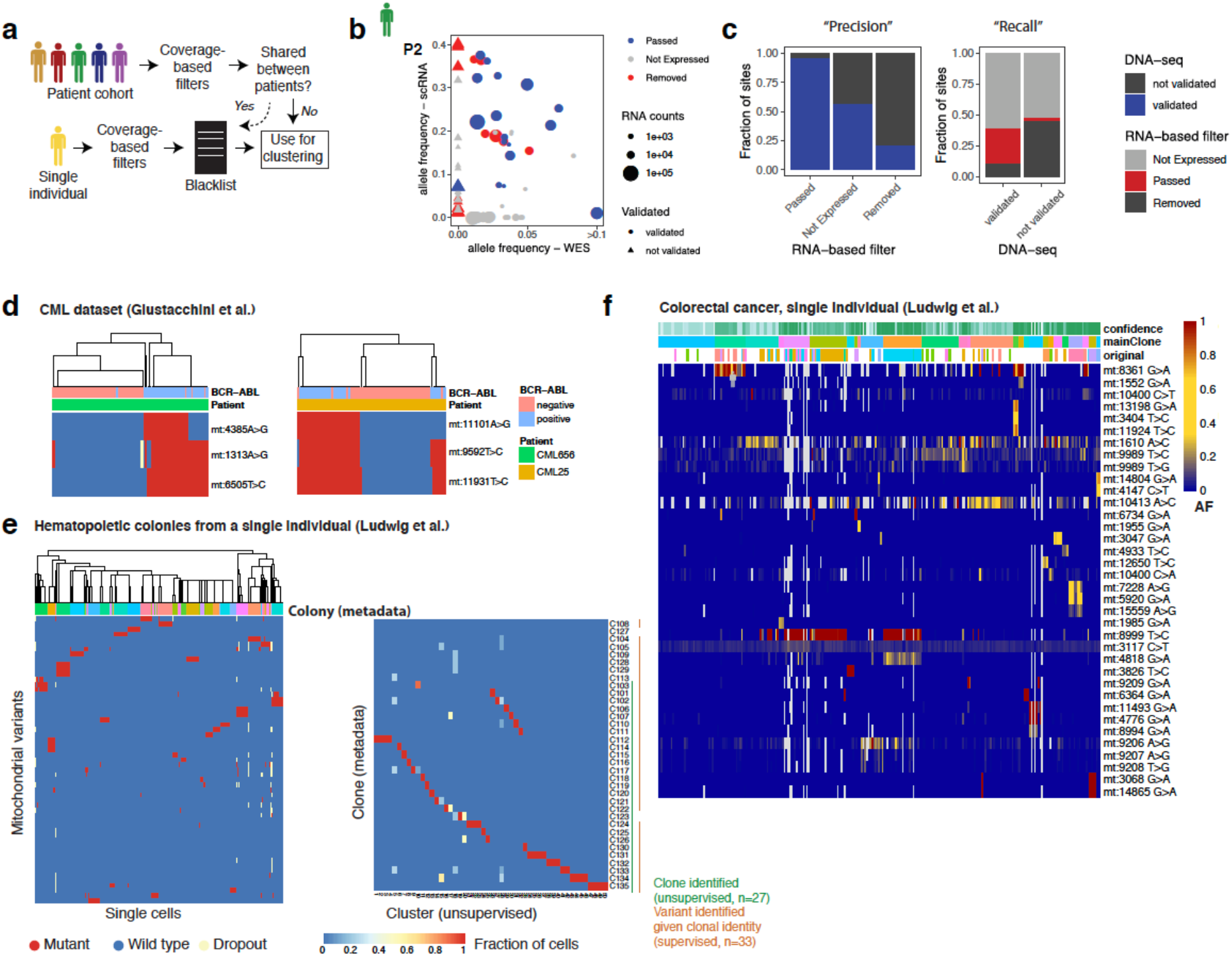
Calling of mitochondrial somatic variants in the absence of a DNA-based reference. See also Figure 2. For an implementation and for reproducing the computations, see the package vignettes of the *mitoClone* package. a. Overview of the computational strategy used. In the case of data from a group of individuals, sites were filtered based on coverage and annotated as ‘mutant’ if a specified fraction of reads deviates from the reference allele. Sites were then excluded as likely RNA editing events if the same mutation was observed in more than one individual. Alternatively, in the case of data from a single individual, similar coverage based filtered were applied and data was then filtered against a blacklist created from a cohort. b. Allele frequencies and coverage of mitochondrial mutations from P2 in single-cell RNA-seq data compared to whole exome sequencing data (WES). Sites with less than 10 reads per cell in RNA-seq were classified as ‘not expressed’. Variants that were observed in WES were classified as ‘validated’ (circles), other variants were classified as ‘not validated’ (triangles). The low correlation between the two datasets is likely due to different starting cell populations (WES: Total bone marrow, single cell RNA-seq: enriched for CD34+ cells), and data are only used for qualitative statements (presence/absence of mutations). c. Bar charts summarizing the classifications from panel b. Left, mutation sites are split by their label based on the *mitoClone* pipeline; right, sites are split by whether they were detected in WES. d. De novo variant calling and clustering of a CML patient dataset. Data from ref. ^67^ were processed and clustered with the *mitoClone* package. The same variant filtering approach used on the patients from our study was used. Thereby, two patients with substantial mitochondrial variability were identified and in both cases clones associated with the BCR-ABL mutation were resolved in an unsupervised manner. The analysis by ref. ^33^ had missed one of these patients, did not achieve an unsupervised separation of BCR-ABL+ and BCR-ABL- cells in either case (Figure 7G in ref. ^33^), and instead relied on stratifying cells by the existing BCR-ABL label (Figure 7J in ref. ^33^). e. De novo variant calling and clustering of single cells from hematopoietic colonies derived from a single individual. Data from Figure 5 of ref. ^33^ were processed and clustered with the *mitoClone* package. Left panel shows unsupervised clustering of mutations identified by the *mitoClone* package, right panel quantitatively compares unsupervised clustering and colony labels. 27 of the colonies were identified in an unsupervised manner. The analysis by ref. ^33^ had identified approximately half that number by unsupervised analyses (their Figure 5E), and using supervised methods identified mitochondrial mutations associated with 33 clones (their Figure 5H). f. De novo variant calling and clustering of single cells from a single colorectal cancer patient. Data from Figure 7 of ref. ^33^ were processed and clustered with the *mitoClone* package. Clustering structure obtained by PhISCS is shown and compared to the clustering presented in ref. ^33^ (row labeled ‘original’), which was based on variant filtering using a DNA-seq based reference. Despite the different filtering approaches, our unsupervised clustering separated the clusters identified by Ludwig et al. and identified additional variability.

**Figure S5.**
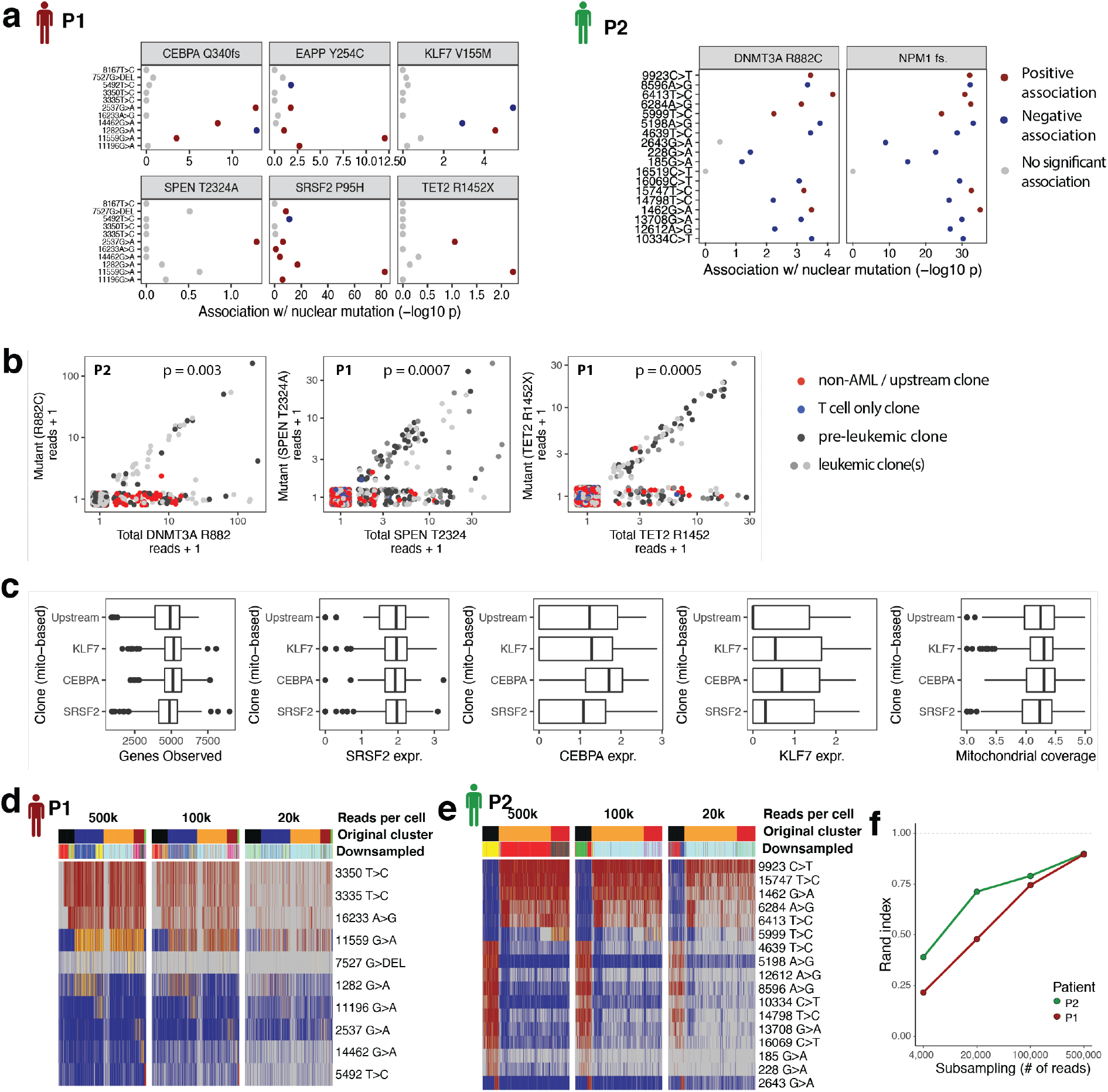
Evaluation of mitochondrial markers for clonal tracking in single-cell RNA-seq data. See also Figure 2. a. Association between various mutations (y axis) and nuclear mutations (panels) across n=1430 cells from P1 or n=1066 cells from P2. P-values are from a two-sided Fisher test. b. Association between the lowly covered mutations in DNMT3A, SPEN and TET2 with clonal identity. Scatter plot depicts total coverage on the site of interest (x axis) and the number of mutant reads (y axis) across n=1066 cells from P2 (left panel) or n=1430 cells from P1 (central and right panel). Note that jitter was added in the x and y direction to avoid overplotting. P values are from a chi-square test comparing a model where the probability of detecting at least one mutant read was modelled as a function of total coverage (null model), or a function of total coverage and identity as a non-AML/upstream clone (alternative model). c. Box plots evaluating the extent to which differences in marker gene expression and library quality affect clonal assignments when using mitochondrial marker mutations; data from P1 is shown. Clone ‘SRSF2’ refers to the pre-leukemic clone from main figure 2 while clones ‘KLF7’ and ‘CEBPA’ refer to the leukemic clones. d. Effect of read depth on mitochondrial clusters. Clusters obtained from mitochondrial sites only were computed at full read depth (row “Original clusters”) and are compared to clusters obtained using the same methodology from data were single cells were down-sampled to read depths of 500k, 100k, or 20k per cell (“Downsampled”). Data from Patient 1 is shown. e. Like panel c, but for patient 2 (P2). f. Original clustering result and down-sampled clustering result are compared quantitatively using the Rand index.

**Figure S6.**
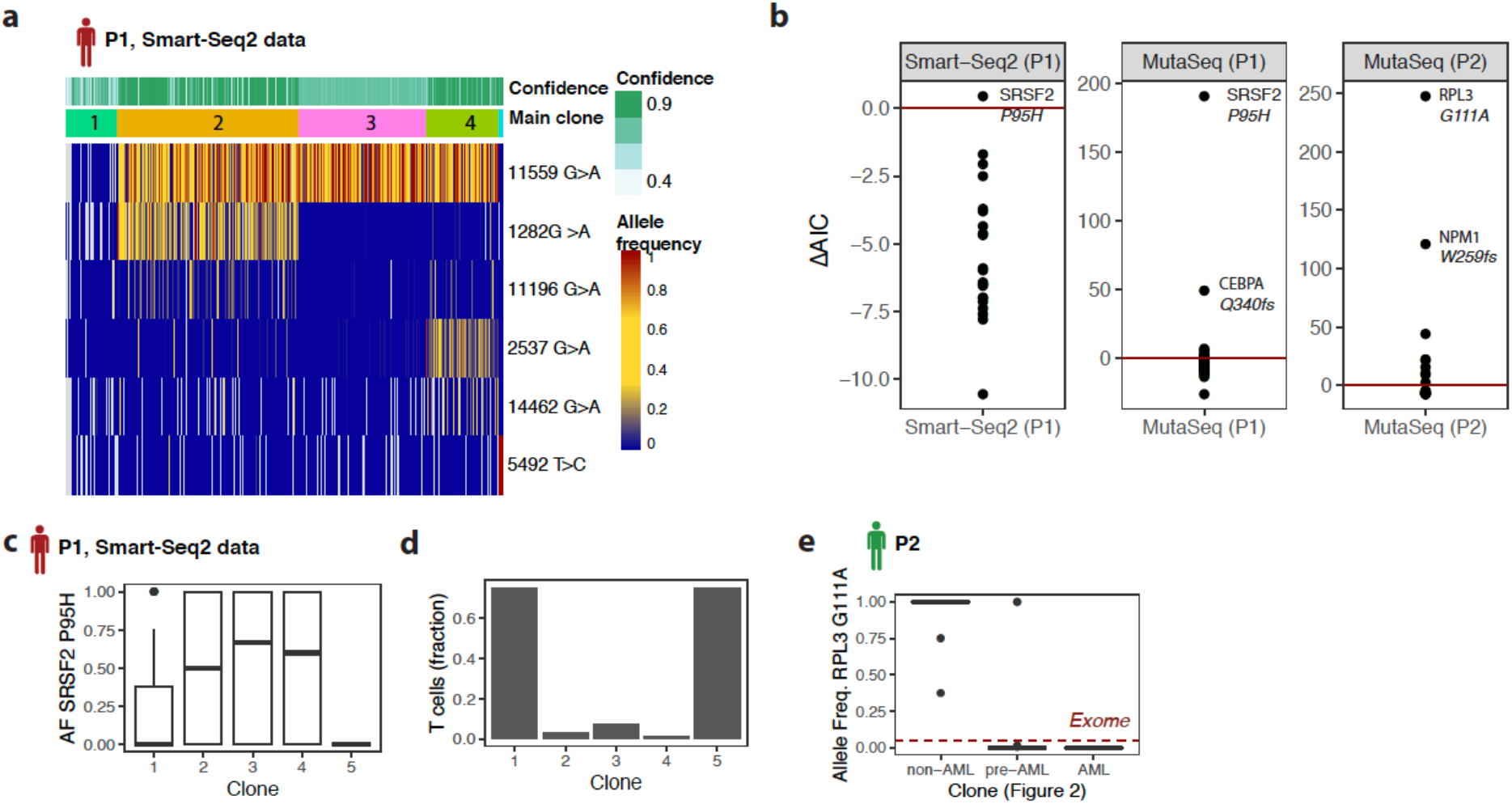
De novo calling and characterization of clones. See also Figure 2. For an implementation and for reproducing the computations, see the package vignettes of the *mitoClone* package. a. Unsupervised clustering of mitochondrial mutations identified from a Smart-seq2 dataset of n=672 cells from patient P1. b. De novo identification of nuclear somatic variants associated with the clonal labels from panel Difference in Aikake’s Information Criterion (AIC) is shown for a comparison between a model where allele frequencies are the same across all cells, and a model where allele frequencies differ between clones. Red line highlights the intercept. See Methods section *Analysis of mitochondrial mutations and reconstruction of clonal hierarchies*. c. Boxplot of single-cell allele frequencies for the SRSF2 P95H mutation summarized between clones (Smart-seq2 data). d. Bar plot of contribution of clones to T cells identified from unsupervised clustering of gene expression data (see also Figure 3). e. Boxplot of single-cell allele frequencies for the RPL3 mutation (COSV53365368) summarized between clones (MutaSeq data). Red dashed line highlights the allele frequency of the mutation identified in exome sequencing.

**Figure S7.**
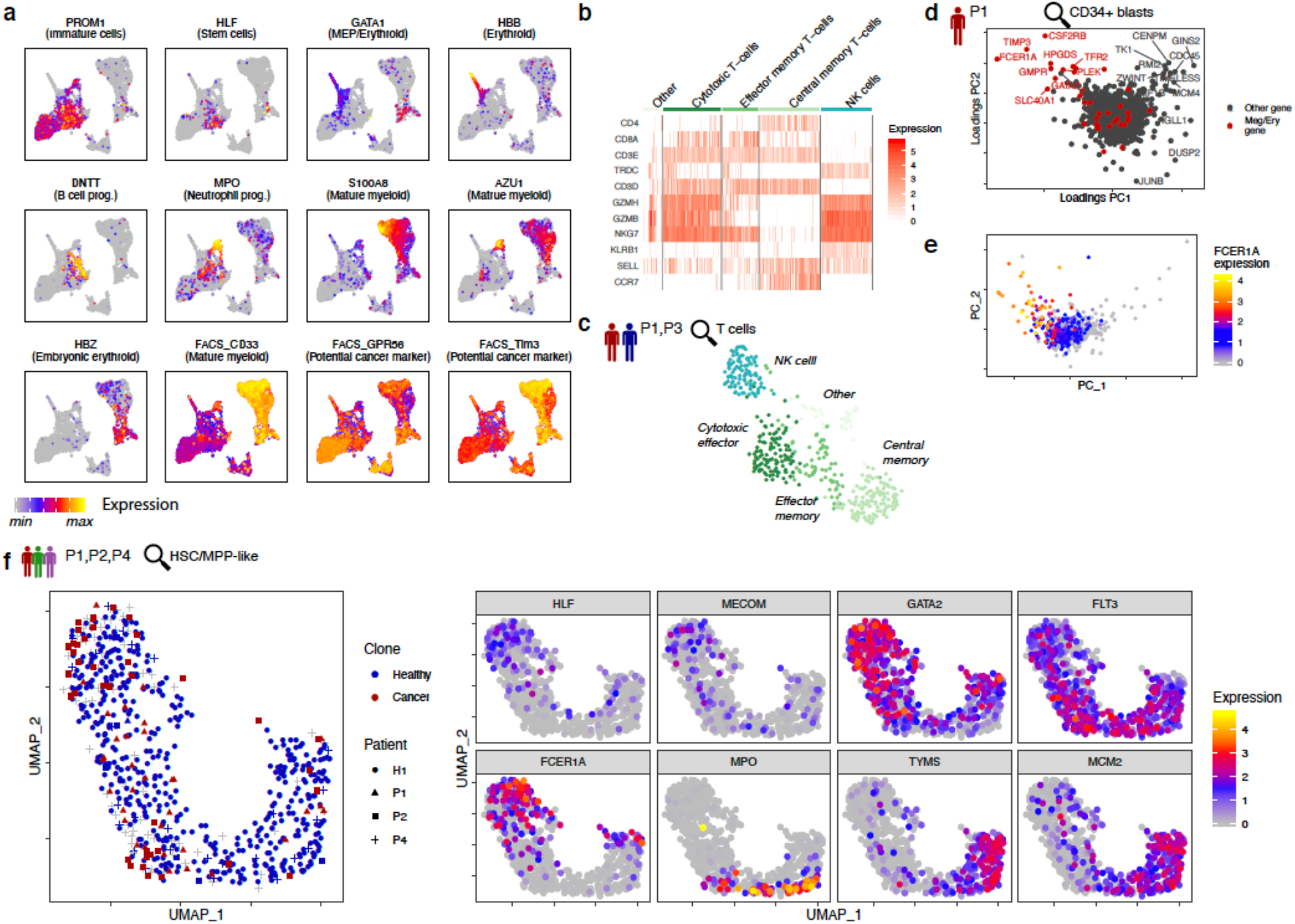
Analysis of single-cell gene expression data. See also Figure 3. a. Expression of marker genes on the uMAP from Figure 3. Top rows: log-normalized mRNA expression. Bottom row (labels preceded FACS_): Logicle transformed FACS index values. b. Heatmap depicting the expression of selected marker genes across T-cells from patients P1 and P3. Columns are cells. c. uMAP representation of all T-cells from patients P1 and P3. Data from these cells only were integrated using MNN^62^ and visualized in two dimensions using uMAP. Patients P2 and P4 were omitted from this analysis due to an insufficient number of T cells. Colors denote cluster identity. d. Loadings plot of a principal component analysis of all CD34+ blasts from Patient P1. Genes associated with erythroid or megakaryocytic priming ^22^ are highlighted in red. e. Scores plot of a principal component analysis of all CD34+ blasts from Patient P1. Expression levels of the FCER1A gene are color-coded. f. uMAP representation of all cells with a healthy HSPC-like gene expression signature from patients P1, P2 and P4. These include both healthy and pre-(leukaemic) clones, see figure 4. Data from these cells only were integrated using MNN^62^ and visualized in two dimensions using uMAP. Left panel: clonal identity is highlighted, using the same strategy as in Figure 4e. Right panels: Point color represents the expression of genes involved in differentiation (MPO for myeloid differentiation, FCER1A for erythroid/megakaryocytic differentiation) and cell cycle (TYMS, MCM2).

**Figure S8.**
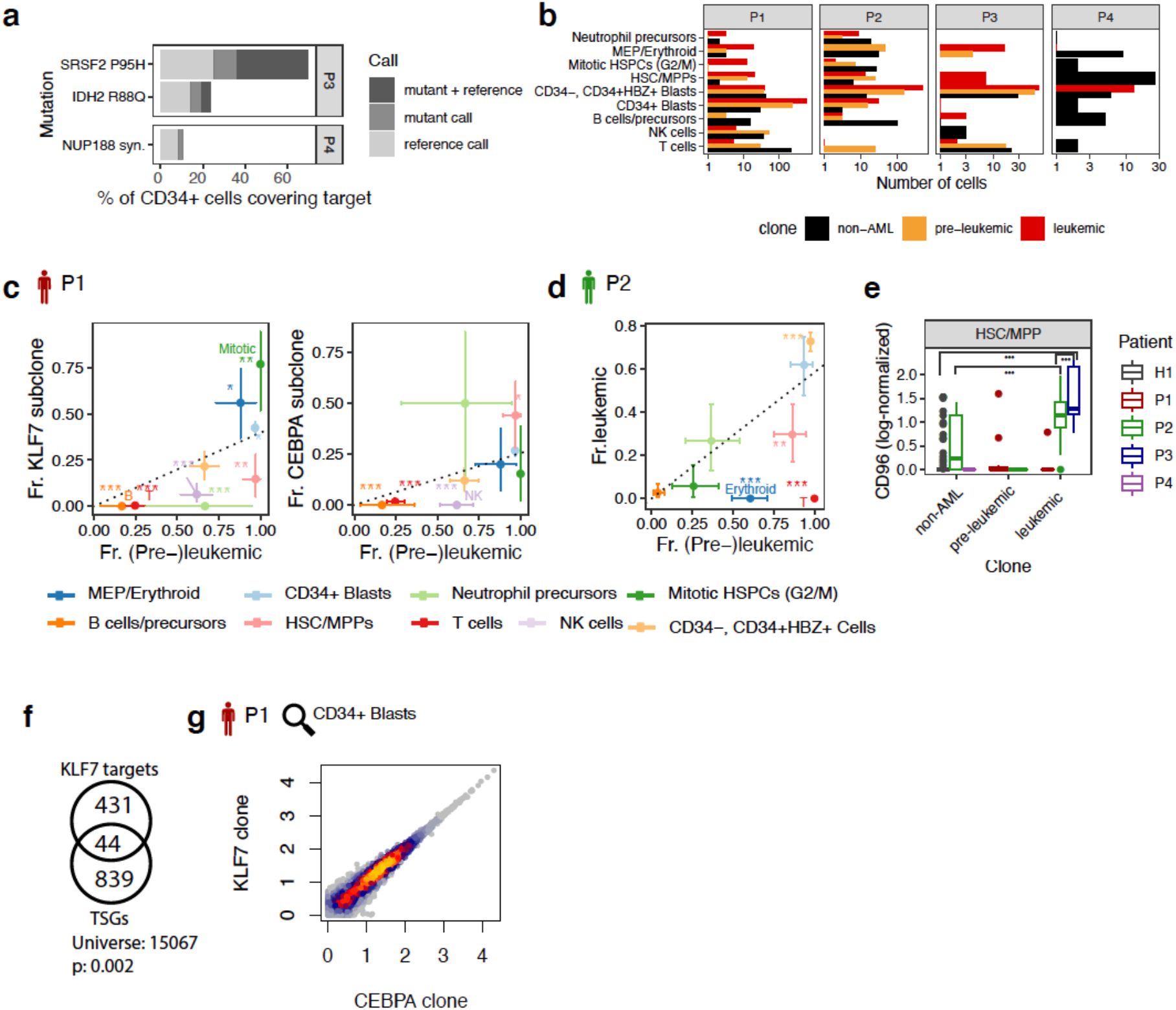
Analysis of single-cell clonal tracking data. See also Figures 4 and 5. a. Bar chart depicting the percentages of cells with coverage of the mutations used for annotating clones in P3 and P4. b. Bar chart depicting the absolute cell numbers of the different clones in the different cell populations. Cells were assigned to clones as in Figure 4e. c. Fraction of (pre-)leukemic cells in relation to the fraction of cells from the sub-clones for P1. Dotted line indicates the mean ratio across all cells, error bars denote 95% confidence intervals from a beta distribution, and asterisk indicate significant deviation from the mean ratio, as follows: *: p < 0.05; **: p < 0.01; ***: p < 0.001. p-values were computed from the quantiles of a beta distribution against the null hypothesis that the true ratio is the mean ratio observed across all cells. Only cells with a confident assignment to clones (likelihood > 0.8, see Methods section ‘Analysis of mitochondrial mutations and reconstruction of clonal hierarchies’) are included. d. Same as Figure S8c investigating instead the second patient (P2). e. Boxplots comparing the normalized, scaled expression levels of CD96 between cells with evidence of originating from the non-leukemic clone(s), and cells with evidence of originating from the leukemic or pre-leukemic clones. Cells were assigned to clones as in Figure 4e. Asterisk denote p-values from a Wilcoxon test for relevant comparisons. ***: p < 0.001 f. Venn diagram depicting the number of open chromatin regions containing a KLF7 binding site, the number of open chromatin regions proximal to tumor suppressor genes from^46^, and their overlap. N=15067 open chromatin regions were identified from single-cell ATAC-seq data of human CD34+ bone marrow cells^45^. g. Scatter plot comparing the log10 mean gene expression levels in CD34+ blasts from the CEBPA-mutated clone, and CD34+ blasts from the KLF7 mutated clone. N=15,451 genes with a mean expression of at least 1 in either population are shown. Color represents point density.

**Figure S9.**
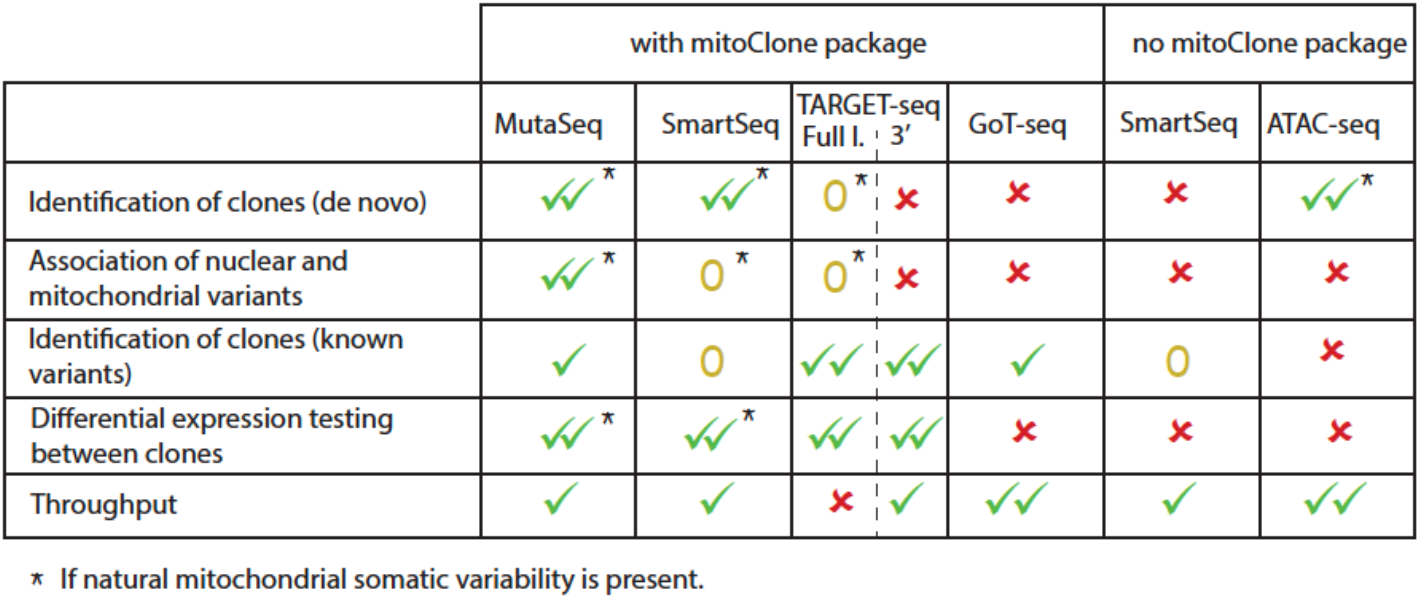
Overview of capabilities of MutaSeq compared to related methods. Full-l.: full-length. SmartSeq2: ref. ^33,54^. TARGET-seq: ref. ^31^. GoT-seq: ref. ^28^. “0” means theoretically possible, but unproven and/or very limited.

**Figure S10.**
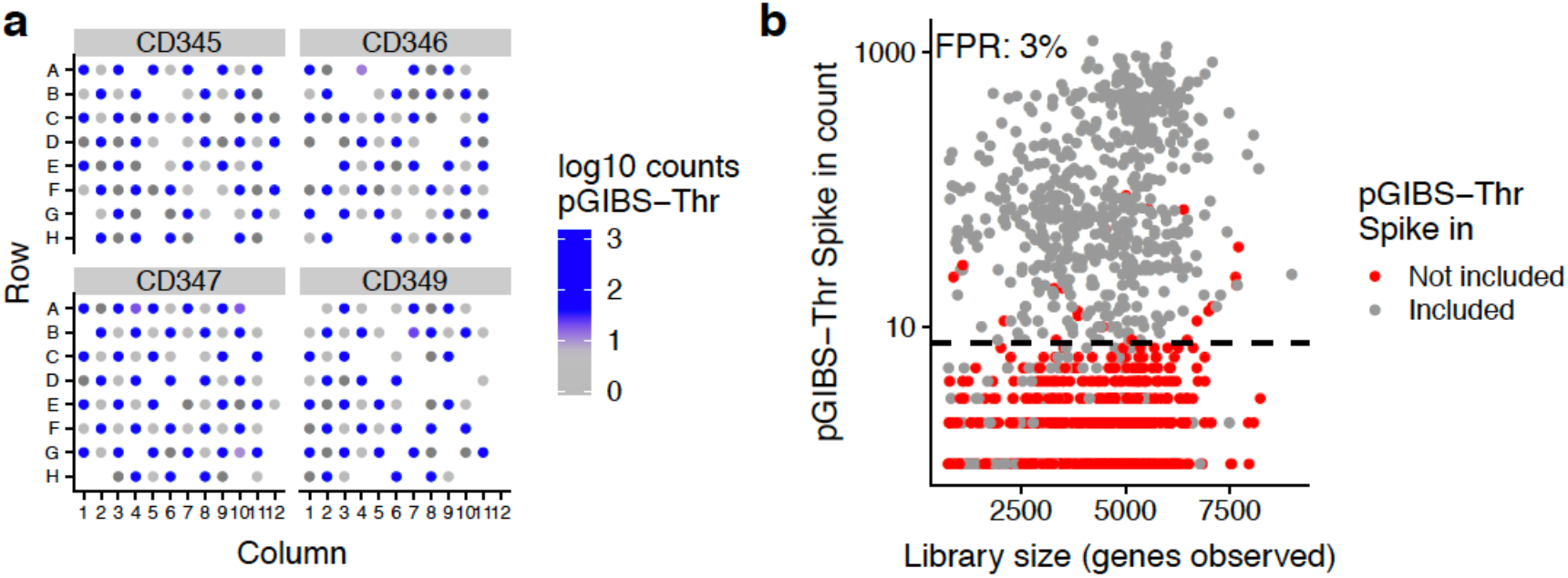
Estimation of MutaSeq’s false positive rate. a. To estimate MutaSeq’s false positive rate, a polyadenylated in vitro transcript, pGIBS-Thr^68^, was added to every second well (A1, A3, B2, B4, etc.) during the P1 experiment. Abundance of the pGIBS-Thr spike-in across wells from four representative plates is shown. b. Estimation of the false positive rate using the pGIBS-Thr spike-in. Dashed bold line indicates the threshold used for classifying a site as dropout.

## Supplementary Materials

Table S1. Exome sequencing results. Columns contain mutation location, reference and alternate alleles, consequence for protein sequence, gene biotype, and allele frequencies in their respective patients’ cancer (af_tumor) and healthy (af_normal) exomes. Patients are separated by tab.

Table S2. Genes with significant (false discovery rate <0.001) overexpression in the various cell populations. For DE testing, MAST^63^ with library quality and patient as covariates was used.

Table S3. Results from differential expression test underlying figures 3g, 5b, 5e Table S4. Antibodies used for flow cytometry.

Table S5. Primers used for targeting genes of interest with MutaSeq.

